# Emergent myxobacterial behaviors arise from reversal suppression induced by kin contacts

**DOI:** 10.1101/2021.04.08.439045

**Authors:** Rajesh Balagam, Pengbo Cao, Govind P. Sah, Zhaoyang Zhang, Kalpana Subedi, Daniel Wall, Oleg A. Igoshin

## Abstract

A wide range of biological systems – from microbial swarms to bird flocks, display emergent behaviors driven by coordinated movement of individuals. To this end, individual organisms interact by recognizing their kin and adjusting their motility based on others around them. However, even in the best-studied systems, the mechanistic basis of the interplay between kin recognition and motility coordination is not understood. Here, using a combination of experiments and mathematical modeling, we uncover the mechanism of an emergent social behavior in *Myxococcus xanthus*. By overexpressing cell surface adhesins, TraA and TraB, involved in kin recognition, large numbers of cells adhere to one another and form organized macroscopic circular aggregates that spin clockwise or counterclockwise. Mechanistically, TraAB adhesion results in sustained cell-cell contacts that trigger cells to suppress cell reversals, and circular aggregates form as the result of cells’ ability to follow their own cellular slime trails. Furthermore, our *in-silico* simulations demonstrate a remarkable ability to predict self-organization patterns when phenotypically distinct strains are mixed. For example, defying naïve expectations, both models and experiments found that strains engineered to overexpress different and incompatible *traAB* allelles nevertheless form mixed circular aggregates. Therefore, this work provides key mechanistic insights into *M. xanthus* social interactions and demonstrates how local cell contacts induce emergent collective behaviors by millions of cells.

**Importance:** In many species, large populations exhibit emergent behaviors whereby all related individuals move in unison. For example, fish in schools can all dart in one direction simultaneously to avoid a predator. Currently, it is impossible to explain how such animals recognize kin through brain cognition and elicit such behaviors at a molecular level. However, microbes also recognize kin and exhibit emergent collective behaviors that are experimentally tractable. Here, using a model social bacterium, we engineer dispersed individuals to organize into synchronized collectives that create emergent patterns. With experimental and mathematical approaches we explain how this occurs at both molecular and population levels. The results demonstrate how the combination of local physical interactions triggers intracellular signaling, which in turn leads to emergent behavior on a population scale.

## Introduction

Living systems display remarkable spatial organization patterns from molecules to cells to populations (1, 2). These patterns are a hallmark of emergent behaviors whereby complex functions arise from simple local interactions. For instance, at the cellular level, we have a relatively good understanding of neuron function, but how a collection of neurons integrates into a functional brain is poorly understood. In other cases, emergent behaviors are driven by the coordinated movement of system parts, as seen in the collective motion of insect swarms or bird flocks (3). Inherent in these processes is the ability of individuals to recognize their kin through brain cognition and adjust their movements relative to others around them. Despite much interest in emergent behaviors, the molecular and mechanistic basis of the interplay between kin recognition and the coordination of movements is poorly understood.

The Gram-negative gliding bacterium *Myxococcus xanthus* is a leading model for studying the molecular basis of microbial kin recognition and, separately, for understanding how cells coordinate their movements (4, 5). These microbes are unusually social and exhibit numerous emergent behaviors. Among these are the formation of traveling wave patterns, termed ripples, in which millions of cells self-organize into periodic, rhythmically moving bands (6–8) and, under starvation conditions, aggregate into multicellular fruiting bodies (9, 10). Notably, these emergent social behaviors form from incredibly diverse microbial populations in soil (11), where *M. xanthus* employs kin discrimination to assemble clonal populations and fruiting bodies (12–14). Central to these social behaviors is the ability of cells to control their direction of movement. These long rod-shaped cells tend to align in dense populations (9, 15) and move along their long axis periodically reversing their motion polarity – head becomes tail and vice versa. Cellular reversals are in turn largely controlled by the Frz chemosensory signal transduction pathway (5). Although much progress has been made in myxobacteria biology, a comprehensive and broadly accepted model that explains their self-organization behaviors and kin discrimination is lacking.

One system *M. xanthus* uses to discriminate against non-kin is based on outer membrane exchange (OME) (13). Here, cells recognize their siblings through cell-cell contacts mediated by a polymorphic cell surface receptor called TraA and its cohort protein TraB. TraAB functions as an adhesin, and cells that express identical TraA receptors adhere to one another by homotypic binding, while cells with divergent receptors do not (16, 17). Following TraA-TraA recognition cells bidirectionally exchange outer membrane proteins and lipids (18). The exchange of diverse cellular cargo, including polymorphic toxins, plays a key role in kin discrimination and facilitating cooperative behaviors (19–21). For these, among other reasons, TraAB-mediated OME in myxobacteria serves as a promising model for emergent behavior control; however, whether and how this actually occurs is unknown.

In this study, we investigate the interplay between TraAB-mediated cellular adhesion and motility coordination. Specifically, elevated cell-cell adhesion forces through overexpression of TraAB drive emergent behaviors involving coordinate movements of thousands to millions of cells. To mechanistically understand this emergent behavior, we recapitulated these behaviors in agent-based simulations that mathematically and mechanistically elucidate how these new behaviors emerge. Specifically, we deduced that an intracellular signal arising from sustained cell-cell contacts, mediated by the TraAB adhesins, results in suppression of cellular reversals and thereby allows millions of cells to move as a uniform collective.

## Results

### TraAB overexpression creates emergent circular aggregate behavior

TraAB cell surface receptors govern allele-specific cell-cell adhesion. When TraAB was overexpressed, cells adhered both end-to-end and side-by-side during shaker flask cultivation (Fig. 1) (16, 19). As myxobacteria are motile on surfaces by adventurous (A) and social (S) gliding motility (22), we sought to understand if TraAB-mediated adhesion affects their collective movements. To clearly assess the impact of cellular adhesion on emergent group behaviors, TraAB adhesin was overproduced from a single copy chromosomal locus in an A^+^S^−^ background (Δ*pilA*), since S-motility promotes extracellular matrix production that complicates analysis. When these cells (hereafter TraAB OE cells) were placed on agar, they displayed an emergent behavior, where thousands of cells self-organized into macroscopic circular aggregates (CAs) (Fig. 1). Initial signs of CAs were easily seen 4 h after cell plating and were prominent by 8-12 h (Fig. S1A and the corresponding Movie S1). Following extended incubation periods, CAs enlarged to millimeters in diameter with each containing millions of cells. In contrast, the parent strain (A^+^S^−^, here referred to as wild type (WT)) does not form CAs. Using a different strain with inducible *traAB* expression, CAs were only seen when cells were grown with an inducer (Fig. S1B). In prior work, smaller and simpler versions of CA-like structures were seen in certain mutant backgrounds and were frequently referred to as swirls (23–26). While CAs superficially resemble precursor aggregates that form into fruiting bodies upon starvation-induced development (27), we emphasize that in our experiemnts TraAB OE cells were grown on nutrient medium that blocks development. Therefore, without engaging in a complex developmental lifecycle, TraAB overexpression provides a simple and tractable system to assess the impacts of local cell-cell interactions on emergent group behaviors.

**Fig. 1.**
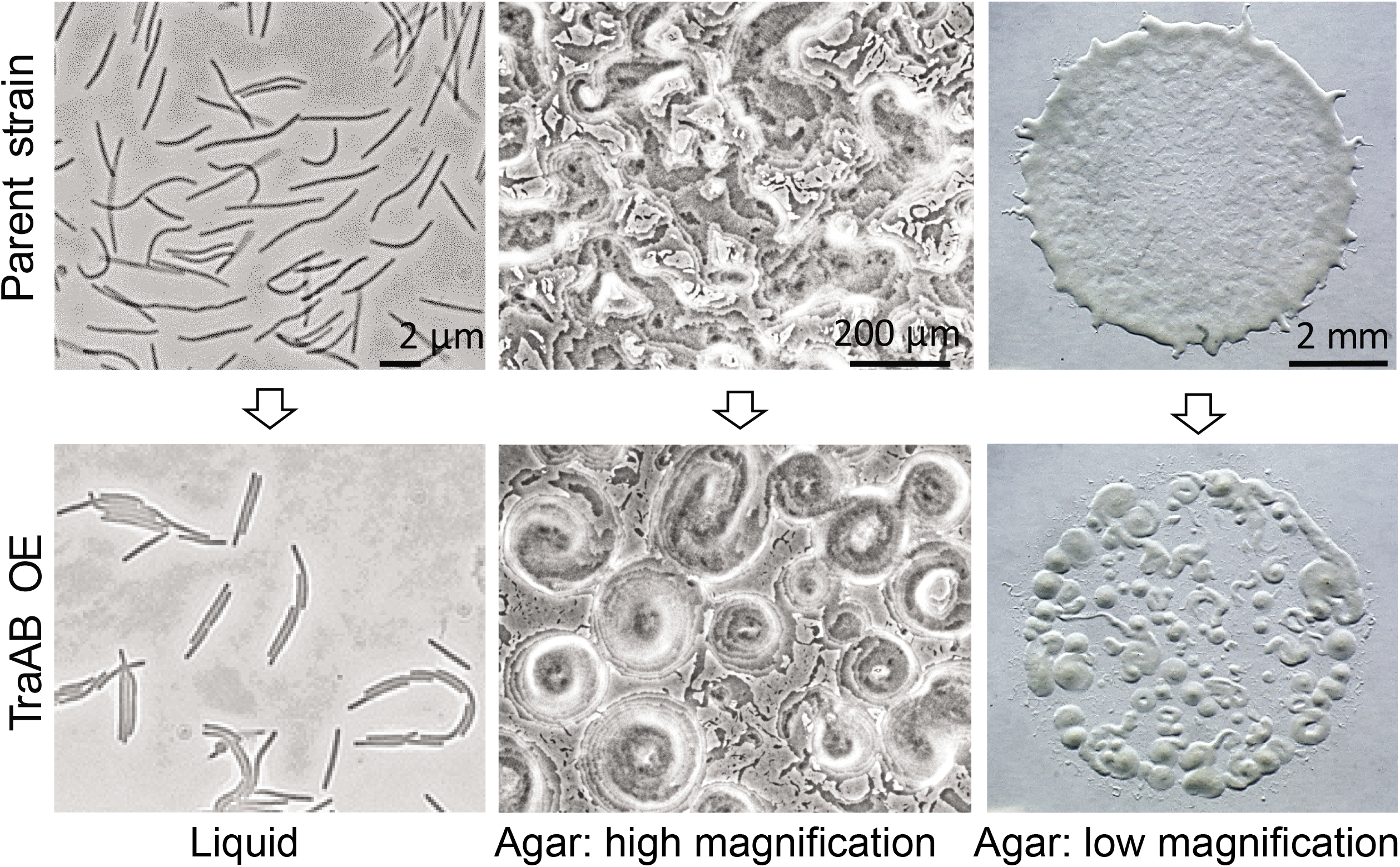
Emergent behavior triggered by TraAB overexpression (OE). Cells adhere from shaker flask growth (left panel), while on agar surfaces motile populations form circular aggregates (CA) when grown on rich media (middle/right panels, 12 h growth). For simplicity, cells only contain one functional gliding motility system.

### The biophysical model reveals CAs only arise from non-reversing agents

To understand mechanistically how CAs emerge in a TraAB OE strain, we attempted to replicate this behavior *in silico* using a biophysical modeling framework that can properly account for forces between cells. To this end, we started with the biophysical model developed by Balagam et al. (28). In this model, to simulate flexible rod-shaped cells, each agent was represented by 7 nodes connected by springs. Agents align with one another on collisions (15) and follow paths left by other agents. These biologically relevant paths are called slime trails composed of poorly characterized material consisting of polysaccharides and lipids that are deposited by gliding *M. xanthus* cells (29–33). Previously, this model was shown to result in CA formation when the slime-trail following was strong and cells did not reverse (15). Notably, physical adhesion between agents was not required in that model of CA formation. However, in light of our experimental findings (Fig. 1), it seemed that TraAB adhesive forces directed CA formation.

To further assess the role of physical adhesion on emergent behavior, we introduced end-to-end and side-by-side adhesion into our model (see Material and Methods for details). The simulation results indicated that the addition of adhesion forces by itself does not promote the formation of CAs when agents have periodic reversals (WT cells reversal period ~8 min (6)). Instead agents self-organized into a network of connected streams (Fig. 2A), with patterns resembling those without adhesion (15). These simulated patterns also resemble experimental observations of the parent strain (Fig. 1, top middle panel). Notably, a further increase in the strength of adhesive forces does not lead to CAs in the population of reversing agents. Instead, excessive adhesion forces exceeding those generated by the agent’s motors resulted in unrealistic bending of agents (Fig. 2B). On the other hand, non-reversing agents in our simulations self-organized into CAs either in the absence (Fig. 2C) or in the presence (Fig. 2D) of adhesion. By varying the reversal frequency of agents we show that CAs only begin to appear when the reversal period exceeds ~70 min, i.e. about 10-fold reversal suppression relative to WT was required for the emergence of CAs (Fig. 2E). Comparing the emergent patterns in Figs. 2C-D, we conclude that in our model side-to-side and end-to-end adhesions by themselves do not significantly affect the emergent patterns. On the other hand, in addition to suppressed reversals, the ability of agents to lay and follow slime trails was critical (Fig. 2F). As groups of cells move unidirectionally along such trails, the natural fluctuation in their trajectories leads these paths to close on themselves so that swirling patterns efficiently reinforce trails to nucleate CAs. As other cells join these swirling paths, CAs grow. Thus, our simulations predict that long reversal periods were necessary for CA formation and, therefore, we predict that TraAB OE cells must somehow alter cellular reversals. However, to date, no connection between TraAB levels and reversal control was known.

**Fig. 2.**
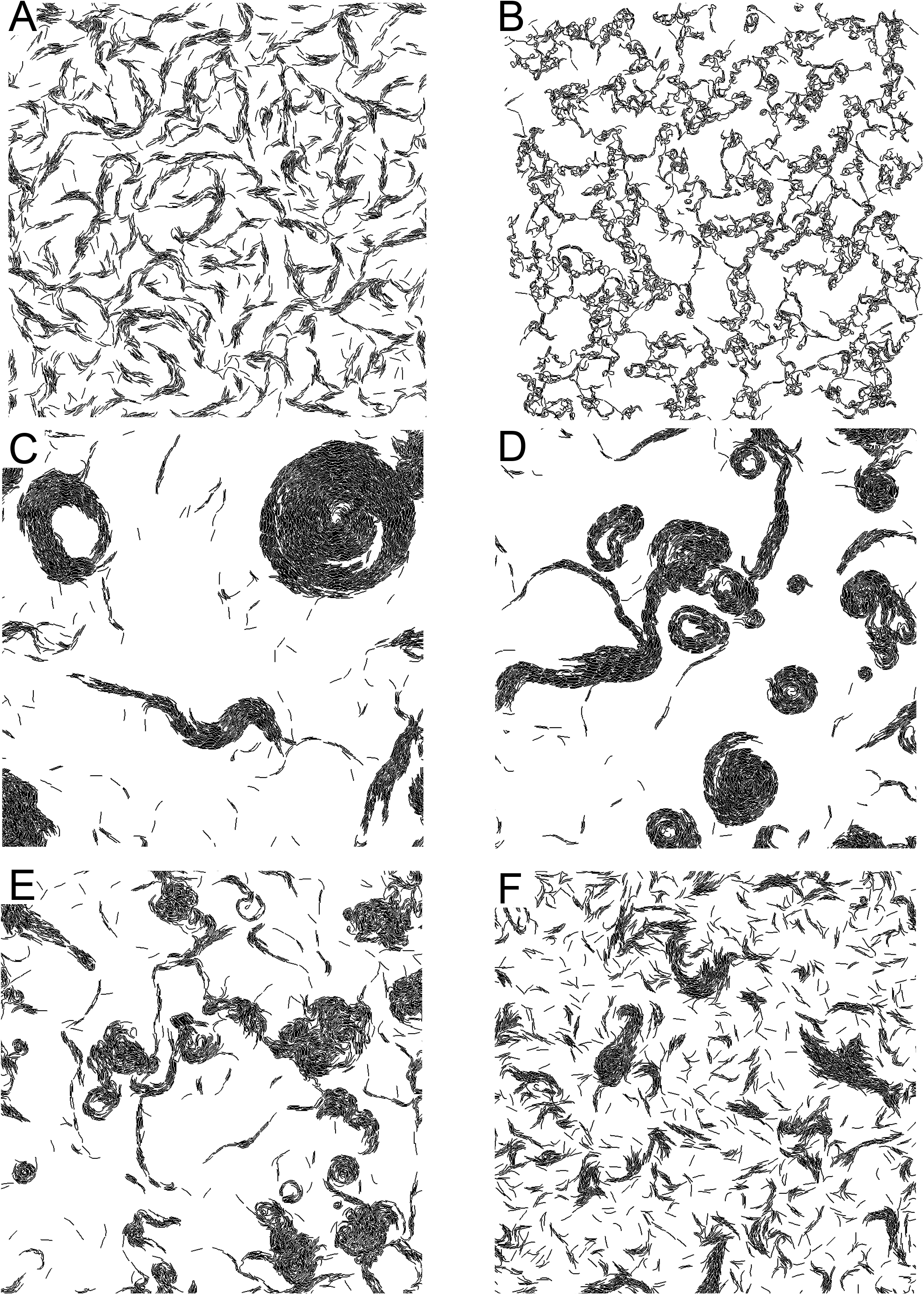
Biophysical model predicts non-reversing and slime-following agents required for CA formation. (A) Reversing agents do not form CAs in the presence of adhesion or (B) with adhesion forces stronger then motor forces. (C) Non-reversing agents form CAs even in the absence and (D) in the presence of adhesion. (E) Reversing agents with long reversing periods (70 min) initiate CA formation. (F) Non-reversing agents without slime-following do not form CAs.

### Cells in CAs suppress reversals

To experimentally test the model prediction, we tracked the movement of single cells within CAs. To this end, a small fraction of TraAB OE cells were fluorescently labeled and mixed with isogenic unlabeled cells (Fig. S2A). Cell movements were recorded by time-lapse microscopy (Movie S2) and the tracks and reversals were quantified as in (9). Figure 3A shows the compiled trajectories of these cells with different (random) colors assigned to individual cells. These trajectories reveal that inside CAs all cells move in the same direction around the center of each aggregate. The CAs themselves rotated in either a clockwise or counterclockwise direction (Fig. S2A and Movie S2). Importantly, when the reversal period was measured for all 443 cells that remained trackable (i.e. in the field of view) for the duration of the movie (60 min), only 12 reversal events were detected. This corresponds to an average frequency of one reversal per cell every ~36 hrs. In other words, cells within CAs did not reverse (Fig. 3B). These results were consistent with our simulation predictions that cell reversals were indeed inhibited within CAs.

**Fig. 3.**
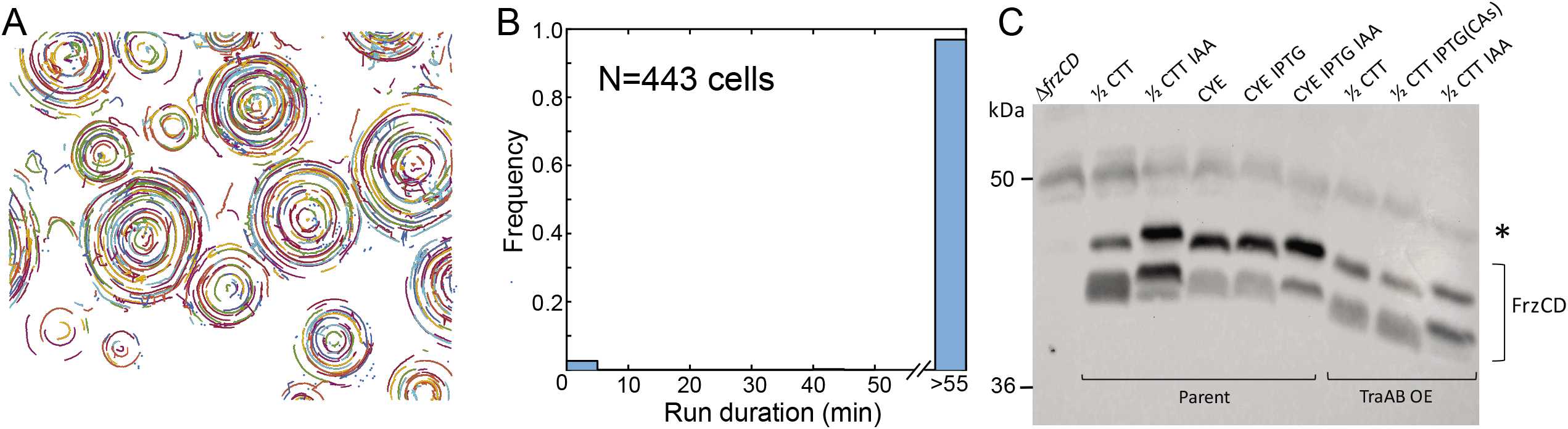
TraAB OE suppresses cell reversals via the Frz chemosensory pathway. (A) Digitally labeled trajectories of marked cells from Movie S2. Each colored curve represents a trajectory of one cell; 443 cells were tracked for the duration of the whole movie (60 min, 1 min intervals). (B) Run duration distribution of the tracked cells: vast majority of cells did not reverse during the observation. (C) CA formation occurs independently of FrzCD methylation changes. A negative control (Δf*rzCD*), parent and TraAB OE strains were harvested from indicated agar plates grown for 24 hr. Only the TraAB OE strain with 2 mM IPTG formed CAs. Representative immunoblot probed with α-FrzCD serum shown. Left, molecular weight standards; *non-specific loading control band.

### Reversal suppression and CAs are dependent on cell-cell adhesion and independent of OME

To determine whether reversal suppression was dependent on cell-cell adhesion or simply due to the TraAB proteins being expressed at elevated levels, the reversal frequencies of isolated cells were also tracked. Here, isolated TraAB OE cells were found not to suppress their reversals compared to controls (Fig. S2B). Additionally, given that TraAB mediates OME, whereby bulk protein and lipid cargo are bidirectional transferred between cells (34), it raises the possibility that reversal suppression, and hence CA formation, was the result of hyper-active OME. To address this possibility, the OmpA domain from TraB was deleted, resulting in a strain producing functional TraAB adhesins, but defective in OME (Fig. S3A, B). Importantly, this strain similarly formed CAs, albeit at reduced levels (Fig. S3C). We conclude that sustained cell-cell contacts mediated by TraAB OE, but not OME, suppresses cell reversals.

### Reversal suppression is required for CA formation

Cellular reversal control in *M. xanthus* is complex. Here a central decision making system is the ‘chemosensory’ signal transduction pathway called Frz (5), which influences the polar localization of the master reversal switch MglA, a small Ras-like GTPase, which in turn determines the polarity of motor function and direction of cell movement. To test the role of reversal suppression in CA formation we first used a chemical inducer (isoamyl alcohol, IAA) of reversals, which acts as a repellant by activating the Frz pathway (35, 36). Here IAA was added at low concentrations to agar media and the behavior of the TraAB OE strain was assessed. Importantly, in a dose-dependent manner CA formation was abolished (Fig. S4A). Secondly, based on our simulations (Fig. 2C) and prior work (24, 26), we confirmed that Frz non-reversing mutants can form CAs in the absence of engineered adhesion (Fig. S4B), although these structures were not as prominent as those in the TraAB OE strain. Additionally, another mutation (Δ*mglC*) that reduces cellular reversal frequencies (25), and apparently functions independently of the Frz pathway (37, 38), also forms CAs (aka swirls) (25), albeit infrequently. Taken together these results support the model that CA formation requires reversal suppression.

Next, we tested whether CA formation, and hence reversal suppression, was signaled through the Frz pathway. As background, similar to other chemosensory pathways in enteric bacteria (39), the Frz pathway contains a methyl-accepting chemotaxis protein (MCP) called FrzCD. However, FrzCD is an atypical MCP that localizes in the cytoplasm and lacks transmembrane and ligand-binding domains (40). Nevertheless, a hallmark of Frz-dependent signaling, similar to other MCPs, are changes in its methylation state as judged by western analysis (41, 42). As previously described (35, 36), in a control treatment with the IAA repellent added to agar media the migration of FrzCD was retarded, indicating an unmethylated state as compared to untreated (½ CTT only) cells (Fig. 4C, upper band). On a nutrient rich agar (CYE), which alters FrzCD methylation and inhibits motility (36), a change in the FrzCD methylation state was also detected as compared to the ½ CTT control. In contrast, when CA formation was induced in the TraAB OE strain by IPTG addition, FrzCD methylation pattern did not change compared to growth in the absence of IPTG (no CAs) or the parent strain grown on ½ CTT (Fig. 4C). However, we note that when SDS-PAGE was conducted under standard conditions, which were not optimized for detecting FrzCD methylation migration differences according to (42), we found minor changes in FrzCD mobility when cells were in CAs (data not shown). Nevertheless, when gel conditions followed the established and optimized protocol for FrzCD (42), we repeatedly found no difference in FrzCD mobility from cells in CAs as compared to controls. Taken together, we conclude that under the optimized assay conditions for detecting FrzCD gel mobility shifts, and hence methylation state, we did not detect appreciable changes when cells were assayed from CAs.

**Fig. 4.**
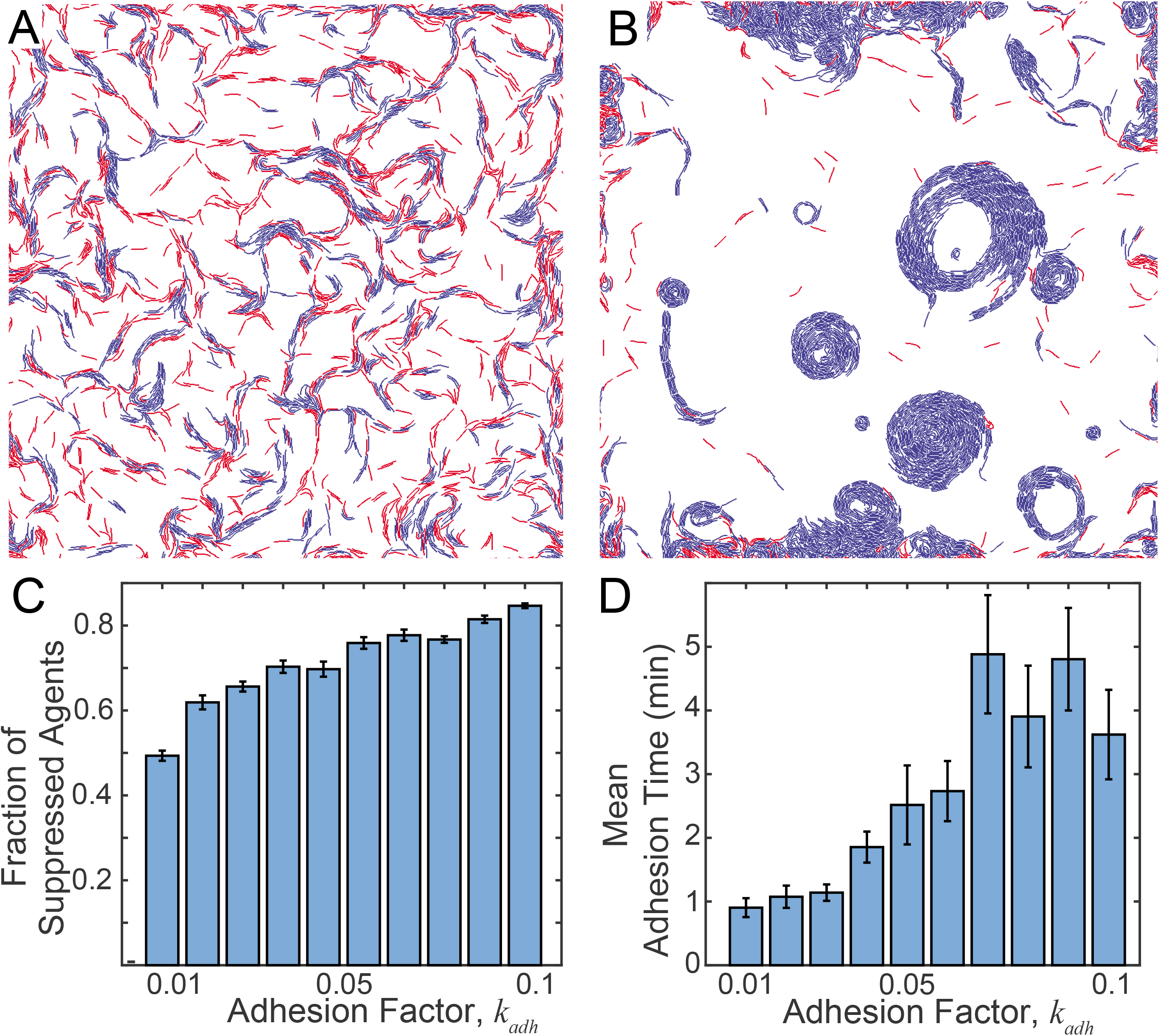
Strong adhesion and cell contact-dependent reversal suppression results in CAs in simulations. (A) Agents with weak adhesion (WT phenotype) show only a small number engaging in reversal suppression (blue) with no emergent behavior. Agents with no reversal suppression are red. (B) Stronger adhesion leads to prevalent reversal suppression (blue) and formation of CAs. (C) Fraction of agents with suppressed reversals as a function of adhesion strength at end of the simulation. (D) Average adhesion bond time at the final 10 min of simulation. The adhesion factor *k_adh_* is defined in SI. WT, *k_adh_* = 0.01; OE, *k_adh_* = 0.1.

### The biophysical model suggests that contact-dependent reversal suppression leads to CA formation

To reconcile the differences in reversal frequencies between cells in CAs (Fig. 3B) and individual cells (Fig. S2B), we hypothesized that sustained cell-cell contacts mediated by TraAB result in an intracellular signal that suppresses cell reversals. Consistent with this model, cell density and contact-dependent signals are known to regulate reversal frequency during development (5, 9, 43, 44), and, additionally, myxobacterial ripples originate from cell contact-dependent reversal modulation (8, 10). To implement this mechanism in our model, we chose a phenomenological approach to simulate contact-dependent reversal suppression inspired by Zhang et al. (10). To this end, at each time step when a given agent was in contact with another agent, its reversal clock was reset backward by a fixed amount. Given that TraAB stimulates both end-to-end and side-to-side adhesion (Fig. 1), we assumed either one or both interactions lead to reversal suppression. We hypothesized that adhesion forces that hold agents together will increase reversal suppression by physically increasing the contact duration. To differentiate WT cell-cell contacts at high-cell densities from those that occur between TraAB OE cells, we introduced a time delay between agent adhesion events and reversal suppression signaling. That delay was set at 5 min to ensure no CAs formed in WT agent simulations as explained below.

In the presence of the signaling delay, with only weak adhesion (representing the parent strain with low TraAB levels; Fig. 4A), agent interactions were short, and reversals were not substantially inhibited, resulting in normal patterns (compare with Fig. 2A). As shown, less than half of the agent contacts lasted long enough to produce reversal suppression. Next, we performed a simulation of agents with stronger and consequently longer adhesion events and as a result, the frequency of reversal suppression was dramatically increased (Fig. 4B). Under these conditions, CAs readily formed, and in agreement with experiments, showed the unidirectional rotation of agents in a clockwise or counter-clockwise direction (Movie S3). This result supports our hypothesis that reversal suppression was necessary for CA formation. Furthermore, our simulations found that when adhesion strength gradually increases from WT to TraAB OE levels the number of agents participating in contact signaling gradually increased (Fig. 4C). However, the effect of the adhesion strength on the duration of adhesion (time before adhesion bond broke) was more dramatic (Fig. 4D). Therefore, to ensure our simulations were consistent with the lack of CAs in the WT strain and based on our findings, we assumed the transient contacts that were shorter than 5 min in the simulations do not suppress reversals. This threshold was important because CAs would form even with weak adhesion in its absence (Fig. S5).

Notably, the behaviors on individual agents in our model match the trends for experimentally tracked cells. With tracking data, we quantified how cell speed and angular speed change as a function of distance to an aggregate center. The experimental results demonstrate that cell speed increases while angular speed decreases as a function of distance from the aggregate center (Fig. S6A, B). Similarly, by quantifying agent speed and angular speed in simulations as a function of distance to the aggregate centers (Fig. S6C, D), we demonstrated that trends were qualitatively consistent with experimental observations (see Fig. S6 legend for details).

### The contact-mediated reversal suppression model accurately predicts emergent patterns of multi-strain mixtures

To further interrogate our model and computationally investigate the interplay between the kin recognition and emergent patterns, we conducted simulations where two types of agents were mixed. In the first simulation, agents overexpressed TraAB receptors of different types (alleles). These receptors do not match and hence the different type agents cannot adhere to each other (16, 17), and reversal suppression only occurs when two agents of the same type engaged in sustained contact. Initially, we hypothesized that differential adhesion would lead to “phase-separation” between agent types, analogous to phase-separation between oil and water. However, in contrast to this prediction, our simulations found that both agent types were mixed within CAs (Fig. 5A). To explain this result, we suggest that reversal suppression and the ability of agents to follow each other’s slime trail overpowered their distinct adhesive forces.

**Fig. 5.**
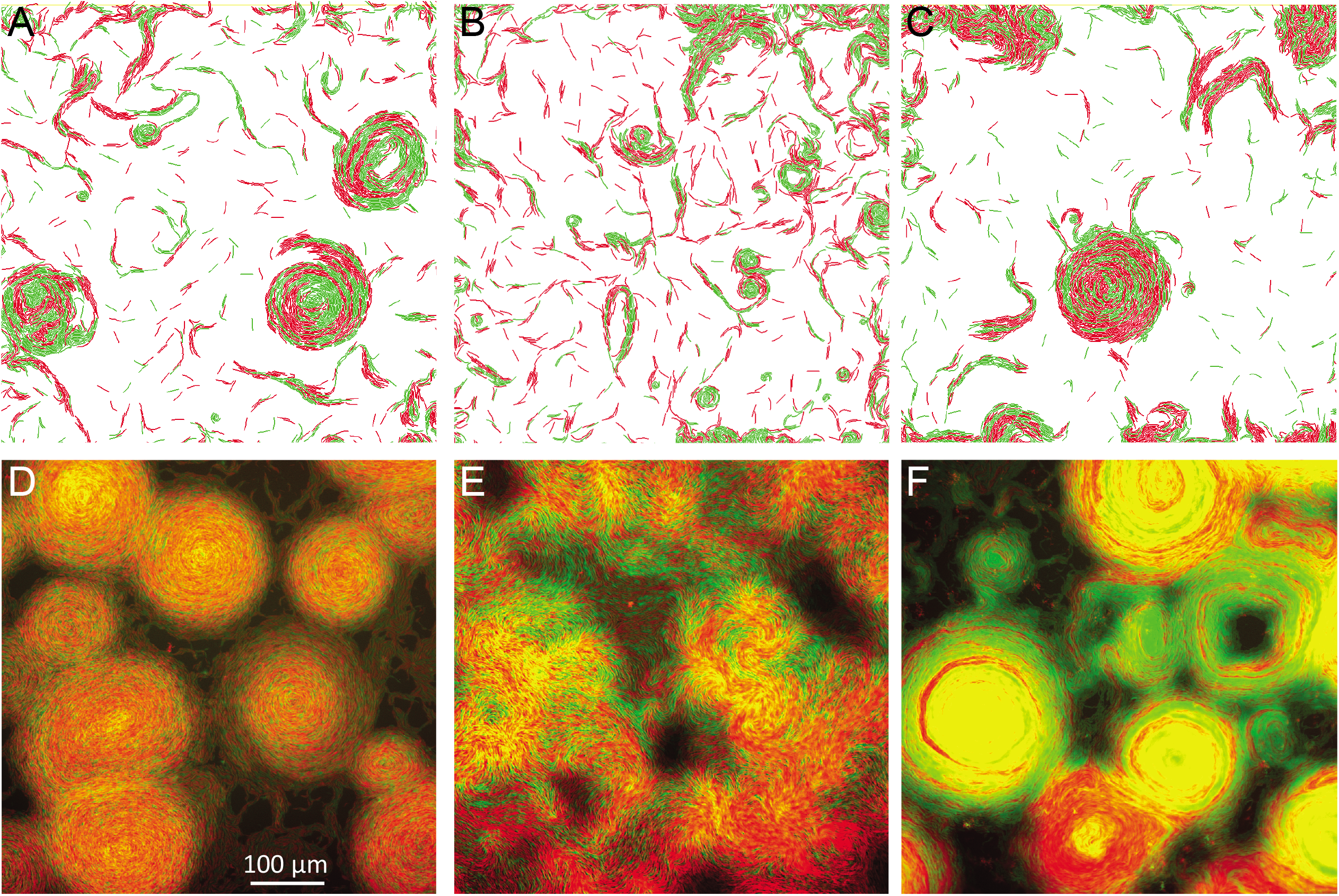
Correlations between simulations and experiments when different combinations of agents or cells were mixed 1:1. (A) Simulation of two different agents (red and green) that adhere to themselves but not each other. (B) Simulation of adhesive agents (TraAB OE, green) mixed with weakly-adhesive agents (WT, red). (C) Simulation of adhesive agents (green) mixed with weakly-adhesive non-reversing agents (red). (D) Experimental mixture of two strains that overexpress different TraA receptors (red and green) that adhere to themselves but not each other. (E) Mixture of TraAB OE strain (green) mixed with a strain that does not adhere (WT, red). (F) Mixture of TraAB OE strain (green) mixed with a non-adhesive non-reversing mutant (red). D-F merged images; see Fig. S7A for single-channel images.

Next, to investigate the impacts of heterogeneous cell-cell adhesion forces across populations, we simulated a TraAB OE (green) mixed with a WT (red) agent. These agents contained different adhesive forces. WT had similar weak adhesions between themselves and with TraAB OE agents (16, 17), and hence they were less susceptible to prolonged cell-cell contacts and reversal suppression. In contrast, TraAB OE agents had strong adhesion among themselves. Interestingly, the simulations showed that the WT agents impeded the formation of CAs by TraAB OE, perhaps by breaking cell-cell adhesions and blocking prolonged contacts that are required for reversal suppression (Fig. 5B, compare to Fig. 4A). Furthermore, to test the role of reversal suppression in CA formation, we conducted a simulation where a TraAB OE agent was mixed with a weakly adhering agent that does not reverse, i.e. a Frz mutant. Strikingly, in this case, the non-reversing and TraAB OE agents formed mixed CAs together (Fig. 5C). This result again demonstrates the key role reversal suppression plays in the emergent CA behavior.

To test our model predictions, we experimentally mixed strains in a manner analogous to simulations. Importantly, for all three strain mixtures, experimental results showed CA patterns or lack thereof, that correlated with all three corresponding simulations (Fig. 5, compare A-C to D-F). Additionally, the degree that strains did or did not mix also correlated well with simulation outcomes, given the latter represents agents in two dimensions while the former shows cells in three dimensions. Specifically, we found that: (i) Introduction of cells with low cell adhesion capabilities (e.g. WT) blocked the emergence of CAs by apparently disrupting prolonged cell-cell adhesions between TraAB OE cells and hence disrupting reversal suppression (Fig. 5E). Moreover, these disruptions were potent because even a minority of such cells, e.g. 7:1 ratio of TraAB OE to WT, reduced CA formation (Fig. S7B). (ii) As found in our simulations and above experiments, reversal suppression played a crucial role, because, in contrast to mixtures with WT cells (Fig. 5E), TraAB OE cells readily formed CAs in 1:1 mixtures with Frz non-reversing mutants, which express TraAB at wild-type levels (Fig. 5F). (iii) When two strains overexpressing incompatible TraA receptors were mixed, they also formed CAs together (Fig. 5D). Therefore, the ability of different strains to strongly adhere to each other was not critical, as long as cell reversals were suppressed, whether by cell-cell adhesion or by *frz* mutations. That is, when divergent populations were mixed, where their cell reversals were suppressed, either by TraAB OE or genetically (*frz*^−^), they readily merged and jointly form CAs by following their reinforced slime trails.

## Discussion

Emergent behaviors transcend the properties of individual components and result in complex functions that are often difficult or impossible to understand mechanistically at a systems level. However, here we investigated a tractable emergent behavior, whereby thousand to millions of cells form spinning CAs. By using experimental and biophysical agent-based modeling, we elucidated the underlying mechanism. Strikingly, our models revealed that the formation of CAs only occurs when cellular reversals are suppressed and cells follow their slime trails, as we previously suggested (15). Experiments confirmed that reversal suppression is required, which is triggered by cell-cell adhesion within dense groups. That is, isolated cells that overexpress TraAB have WT reversal frequencies and necessarily are not constituents of CAs. Using these observations, we hypothesized that reversals were suppressed by long-lasting cell contacts that adhesins stabilized. This model is supported by several experimental findings, including that TraAB OE cells do not reverse within CAs and when reversals are induced by IAA addition CAs cannot form. Secondly, CA formation is phenocopied to some extent by mutants (e.g. *frz*) that are blocked in reversals. Third, our model not only explained the differences in patterns between WT and TraAB OE cells but also qualitatively matches how actual cellular linear and angular speeds change within CAs. Finally, our model accurately predicted emergent patterns when strains with distinct behaviors were mixed.

Central to our model, the formation of CAs only requires cells to lay and follow slime trails (29–31), and adhesion to stabilize cell-cell contacts, thereby leading to reversal suppression. Strikingly, however, for non-reversing agents (or strains), the requirement of adhesion forces can largely be bypassed. This conclusion is supported by the observation that Frz non-reversing mutants form detectable amounts of CAs (Fig. S4B and (24, 26)) and that Δ*mglC* mutation that reduces cellular reversal frequencies also induces similar patterns (25). Thus TraAB-driven cell adhesion is primarily required for reversal suppression rather than for the formation of CAs *per se*. In contrast, another theoretical study showed that non-reversing agents could also form CAs by instead invoking a short-range active guiding mechanism (45). In this model, agents do not follow slime trails, but instead, generate active guiding forces that allow the lagging agent to seek and maintain a constant distance from the leading agent. This active guiding force is assumed to arise from physical adhesion and/or attraction between cell poles, which could be generated by polar type IV pili. Importantly, these models make different predictions on CA dynamics. In one case CAs rotate as rigid bodies (45), whereas CAs based on slime trail following (15) showed that despite the increase in speed there is a decrease in angular velocity the farther agents were from the aggregate center. These patterns of cell speed and angular velocity from experiments qualitatively match our model predictions (Fig. S6) and do not match the predictions of (45). However, since simulations were performed in a single agent layer, which contrast with multilayer cell experiments, the size of simulated CAs remains smaller and thus no quantitative agreement between simulations and experiments is expected. In this context, it is foreseeable that cell adhesion between cell layers further stabilizes CAs and allows them to grow much larger. However, conducting such simulations requires alternative modeling formalism and beyond the scope of this work.

Laboratory competition experiments and characterization of cells from environmentally-derived fruiting bodies reveal that robust kin discrimination systems lead to near homogenous segregation of kin groups from diverse populations (12, 14, 46). OME, mediated by TraAB, plays a central role in these processes by exchanging large suites of polymorphic toxins (13, 20). This ensures that only close kin survive these social encounters because they contain cognate immunity proteins. Here, we found that overexpression of TraAB from kin cells results in the formation of organized social groups that move in synchrony. However, surprisingly, *in silico* and experimental overexpression of divergent TraA recognition receptors, thus representing distinct kin groups, or genetic suppression of reversals by *frz* mutations, resulted in mixed populations within CAs (Fig. 5). Importantly, however, these mixed laboratory groups were between engineered strains derived from the same parent, and hence they were socially compatible because they contained reciprocal immunity to OME toxins as well as type VI secretion system toxins (14). In other words, consistent with ecological findings from fruiting bodies (12), we do not expect mixed CA formation between divergent *M. xanthus* strains that antagonize one another, and thus serving as a barrier to social cooperation (14).

Our findings on CA formation also provide insight into the natural emergent behavior of development. That is, during starvation-induced development cells form spherical fruiting bodies; a process that requires the Frz pathway and reversal suppression (5, 9). Although much is known about development (47), how fruiting bodies emerge remains poorly understood. In light of our findings, we suggest that during development cells increase their adhesiveness, perhaps mediated by C-signaling (48–50), which results in sustained cell-cell contacts, and hence reversal suppression, which similarly is critical for fruiting body formation (9, 44). In a second developmental behavior, cell collision-induced reversals are known to trigger rippling (8, 10). Here we suggest that these collisions could break long-standing cell-cell contacts of aligned groups of cells, thus disrupting their reversal-suppression and triggering reversals. Future studies need to investigate how cell-cell adhesion and sustained cell contacts might change during development and the roles they play during fruiting body morphogenesis and rippling.

Central to the sociality of *M. xanthus* is the control of their cellular reversals that coordinates their multicellular behaviors. Although significant progress has been made in understanding the molecular regulation of reversals (5, 37, 51), major knowledge gaps remain. Here we show that engineered sustained cell-cell contacts suppress cellular reversals. Our findings indicate that the methylation state of the FrzCD MCP is not altered, suggesting Frz mediated adaptation is not involved in reversal suppression. Nevertheless, given that the Frz system plays a major role in reversal control and yet has no known ligand binding domain, our findings do not exclude the possibility that a downstream component, such as FrzE or FrzZ, senses and signals sustained cell-cell contacts. Alternatively, reversal suppression could occur independently of Frz. For example, other systems that regulate reversals include the Dif chemosensory pathway as well as the MglC, PlpA and PixA proteins (25, 37, 38, 41, 52, 53). Additionally, there are undiscovered pathways that suppress reversals as exemplified by the EPS (extracellular polysaccharides) signal (54). Finally, in an alternative scenario, TraAB-dependent cell-cell adhesion could mechanically block the A-motility motor from physically switching cell poles and hence suppress cellular reversals. Consistent with this model, TraAB and the A-motility motor reside in the cell envelop and are mobile macromolecular complexes frequently found at the poles (16, 17, 22, 51).

In summary, our approach provides a roadmap for how strain engineering and modeling helped to elucidate mechanistic insights into an emergent behavior that arises from cell reversal control. These insights are also likely relevant for the natural emergent behavior of fruiting body development. By extension, in other biological systems and model organisms, seemingly complex emergent behaviors, can be broken down and tackled by using a combination of modeling and simplified experimental manipulations to uncover their mechanisms of action.

## Materials and methods

### Bacterial strains and growth conditions

All strains used in this study are listed in Table 1. *M. xanthus* cells were routinely grown in CTT medium (1% [w/v] Casitone; 10 mM Tris-HCl, pH 7.6; 1 mM KH_2_PO_4_; 8 mM MgSO4) in the dark at 33 °C. For a nutrient rich medium CYE was used (1% Casitone, 0.5% yeast extract, 8 mM MgSO4, 10 mM MOPS, pH 7.6). 1.5% (w/v) agar was added to the medium to make plates. To prepare agarose pads for microscopy, Casitone was reduced to 0.2% (w/v), and 1% (w/v) agarose was added to the medium. *E. coli* strains were routinely cultured in LB medium at 37 °C. As needed for antibiotic selection or protein induction, 50 μg ml^−1^ of kanamycin (Km), 10 μg ml^−1^ oxytetracycline, or 1 or 2 mM IPTG was added to media.

**Table 1.**
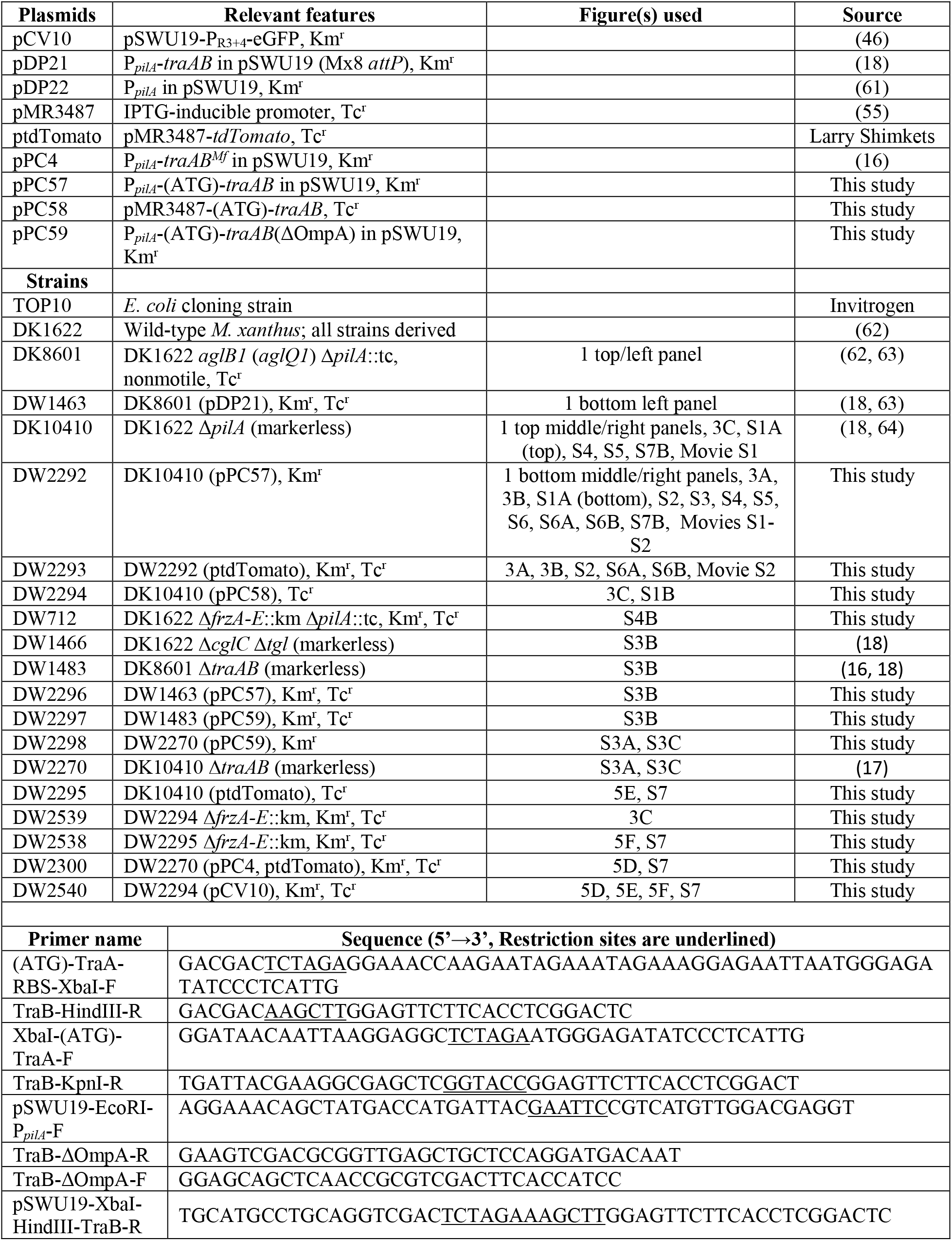
Plasmids, strains and primers used in this study

### Plasmid and strain construction

All plasmids and primers are listed in Table 1. To maximize the expression of TraAB adhesin, pPC57 was constructed where the native GTG start codon of *traA* was changed to ATG and TraAB expression is driven by a heterologous *pilA* promoter (P*_pilA_*). This site-directed mutagenesis was done by using primers containing the desired mutation, and the amplified *traAB* fragments were ligated into pDP22 (linearized with XbaI and HindIII) with T4 DNA ligase. To achieve inducible overexpression of TraAB (pPC58), *traAB* fragments were PCR amplified and then ligated into pMR3487 (linearized with XbaI and KpnI) through Gibson Assembly (New England Biolabs). To create pPC59, primers were designed to amplify fragments of *traAB* and omit the region encoding for OmpA, and the resulting fragments were ligated into XbaI and HindIII digested pDP22 through Gibson Assembly. Plasmid construction was done in *E. coli* TOP10. All plasmids were verified by PCR, restriction enzyme digestion, and if necessary, by DNA sequencing. To construct *M. xanthus* strains, plasmid or chromosomal DNA was electroporated into cells and integrated into the chromosome by site-specific or homologous recombination. For pSWU19 derived plasmids integration occurs at the Mx8 attachment site, while pMR3487 recombines at another site and expression is induced with IPTG (55).

### Aggregate formation

*M. xanthus* cells were grown to logarithmic growth phase in CTT, washed with TPM buffer (CTT without Casitone), and resuspended to the calculated density of 5 × 10^8^ cells per mL. 5 μL of cell suspension was then spotted onto ½ CTT (CTT medium with 0.5% Casitone) agar plates supplemented with 2 mM CaCl_2_. In some cases, different strains were mixed at desired ratios before spotting. Spots were air-dried and plates were then incubated at 33 °C overnight before imaging. When necessary IPTG was added during liquid and plate growth. To assess the impacts of cellular reversals on CA formation, isoamyl alcohol (IAA) was supplemented to agar media at indicated concentrations.

### Microscopy

CA formation on agar plates was imaged using a Nikon E800 phase-contrast microscope (10× phase-contrast objective lens coupled to a Hamamatsu CCD camera and Image-Pro Plus software), or an Olympus IX83 inverted microscope (10× objective lens coupled to a ORCA-Flash 4.0 LT sCMOS camera and cellSens software), or an Olympus SZX10 stereomicroscope (low magnification coupled to a digital imaging system). To track isolated cell reversals, cells were mounted on an agarose pad and imaged with a 20× phase-contrast objective lens. Fluorescence microscopy was used to track individual cells within CAs with a 10× lens objective and a Texas Red filter set. Cell-cell adhesion was imaged directly from overnight cultures mounted on glass slides with a 100× oil immersion objective lens.

### Immunoblot

To optimize separation of different FrzCD isoforms, SDS-PAGE was done as essentially described by McClearly et al (42). Briefly, equal amounts of cell extract were separated on a 14 cm resolving gel consisting of 11.56% acrylamide, 0.08% bis, 380 mM Tris pH-8.6, 0.1% SDS, 0.1% ammonium per sulfate and 0.04% TEMED. The stacking gel consisted of 3.9% acrylamide, 0.06% bis, 125 mM Tris pH-6.8, 0.1% SDS, 0.1% ammonium per sulfate and 0.01% TEMED. To remove non-specific binding, the rabbit α-FrzCD serum was first pre-absorbed against a blot from a Δ*frzCD* strain and then used at a 1:15,000 dilution on experimental blots. For detection, HRP-conjugate goat-anti-rabbit secondary antibody was used (1:15,000 dilution, Pierce) and developed with SuperSignal West Pico Plus chemiluminescent substrate (Thermo Scientific).

### The Agent-Based-Simulation framework

The simulation model framework is adapted from our previous work (15, 28). A brief description of the previous model, simulation framework as well as the new changes introduced in the framework are presented below. All of the parameters are summarized in Table 2. Each agent is represented as a connected string of N (= 7) circular nodes with a total cell length *L* (= 6 μm) and width *w* (= 0.5 μm) (see figure S1 in (28) and additional details in (15)). Neighboring circular nodes are kept at a fixed distance apart by (N-1) rectangular spacers. Neighboring circular nodes and rectangular spacers are connected by linear (spring constant, *k_l_*) and angular (spring constant, *k_b_*) springs. Linear springs here resist elongation and compression of cell nodes. The linear spring constant is managed by the model engine to keep the agent length constant. Angular springs resist bending from straight-line configuration to simulate elastic bending behavior of *M. xanthus* cells.

**Table 2.**
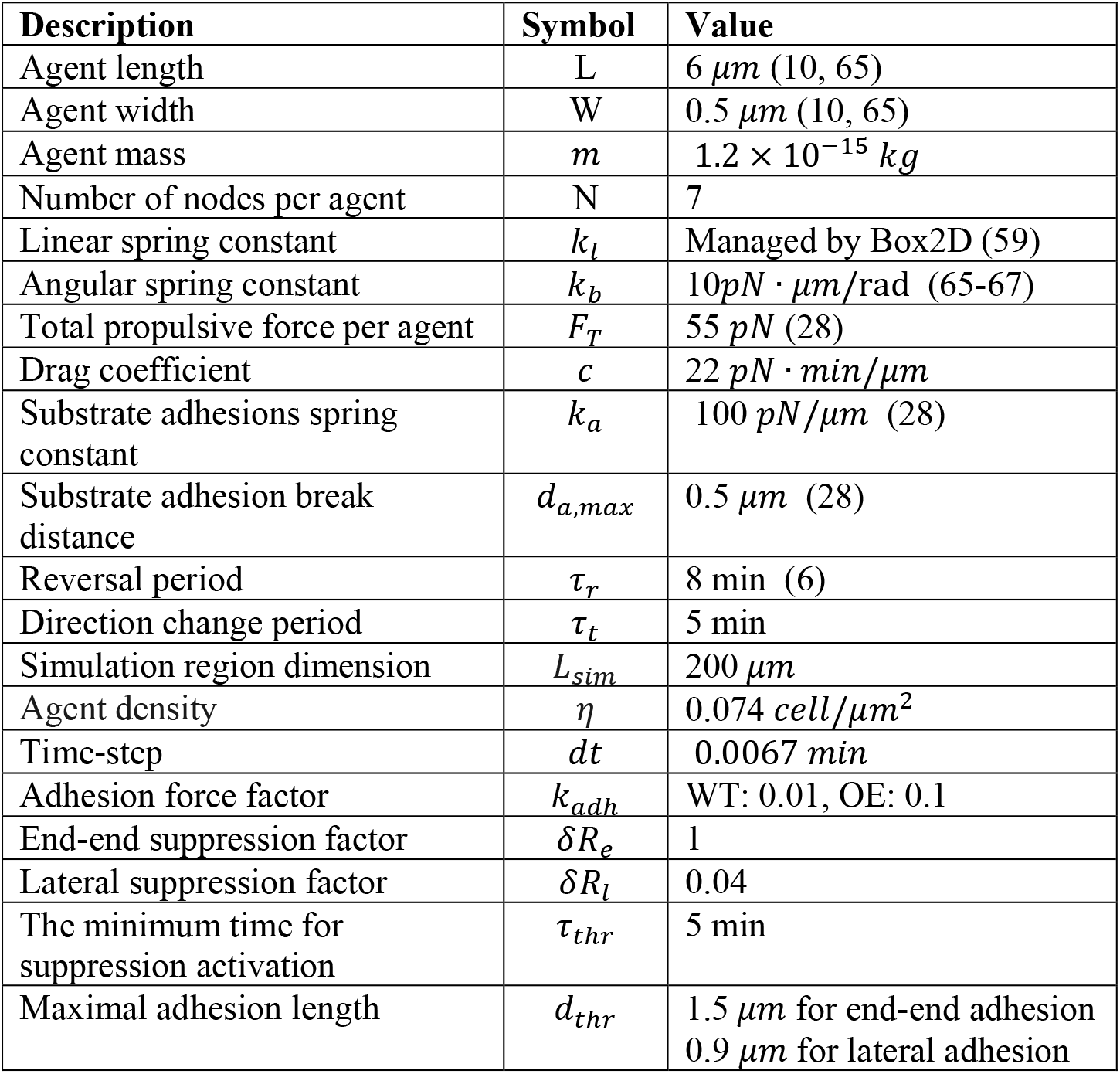
Simulation parameters

Each agent moves forward by the propulsive forces. Since the experiments are performed with the cells lacking S-motility, we only implement gliding (A) motility of *M. xanthus* cells based on distributed force generation model (22, 56–58). At each node *i* a propulsion force (***F**_p,i_* = *F_T_*/(*N* – 1), *F_T_* is the total propulsive force) is applied in the current travel direction towards the neighboring node. Viscous drag forces (***F**_d_*) arising from the surrounding fluid/slime act on nodes opposing their movement with the force proportional to the velocity or each node with proportionality coefficient *c* (drag coefficient).

Agent movement is affected by collisions, periodic reversals, random turns, and slime-trail-following by agents. Collisions in our model are resolved by applying repulsion forces on nodes that keep agents from overlapping. Further, adhesive attachments between the agent and the underlying cell substrate (based on focal adhesion model of gliding motility in *M. xanthus* (57)) at each node resist lateral displacement of the nodes during collisions with other cells. These attachments are modeled as linear springs (spring constant, *k_a_*) and are detached at a threshold distance *d_a,max_*. For each agent, the first and last nodes in the current cell travel direction are designated as head and tail nodes respectively. Periodic reversals in our model are introduced by switching the roles of head and tail nodes and reversing the propulsive force direction at the inner nodes. Reversals in agents are triggered asynchronously by an internal timer expiring at the end of the reversal period (*τ_r_*) after which the timer is reset to zero. *M. xanthus* cells exhibit random turns during movement on solid surfaces (28). These random turns are added to the model by changing the direction of the propulsive force on the head node of the agent by 90° (either clockwise or anti-clockwise chosen randomly) for a fixed amount of time (1 min) at regular time intervals (*τ_t_*) triggered another internal timer. Slime-trail-following by *M. xanthus* cells is a known phenomenon (29), in which cells leave a slime trail on the substrate and other cells crossing these trails later start following them. We added slime-trail-following of agents in our model using a phenomenological approach where we gradually change the direction of propulsive force (***F**_p_*) on the head node of the agent parallel to the direction of slime trail (*ê_s_*) it is currently crossing. (See (15) for implementation details of slime-trail-following mechanism in our model).

### Cell adhesion

To simulate adhesive interactions between agents, we apply lateral adhesive forces (*F_adh_*) on nodes of neighboring agents if the two nodes are closer than a specific threshold distance. In the simulation, we include end-end adhesion where one agent’s head node is attached to another agent’s tail node and lateral adhesion where an agent is attached to a nearby agent side by side. The threshold distance for lateral adhesion *d_thr_* = 0.9 μm and for end-end adhesion *d_thr_* = 1.5μm. This is because we assume the cell wall/membrane can be stretched more along the long axis.

We use the following equation to calculate cell adhesion force:

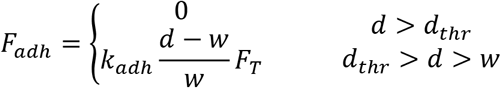

Here, *d* is the distance between neighboring nodes, *w* is the width of cells, *k_adh_* is the adhesion force factor describing the ratio of the maximal adhesive force to the total propulsive force of the agent *F_T_*. For OE cells, *k_adh_* = 0.1 for WT cells *k_adh_* = 0.01. These adhesive forces are applied on each node in the direction towards the neighbor node center.

### Reversal suppression induced by cell contacts

In the model, cell reversal is controlled by a reversal clock in the agent. If the reversal clock records a time longer than the chosen reversal period, the reversing happens and the reversal clock is reset to 0. In this work, we assume if the adhesion lasts longer than a threshold time (*τ_thr_*, set to be 5 min unless indicated otherwise), agents suppress their reversals. We set the threshold to be 5 minutes. When suppression of reversal happens, the reversal clock is slowed down or even turned back for every time step that agents remain in contact past the threshold. For each agent, we calculate the total suppression from end-end pairs and lateral suppression contacts, i.e.:

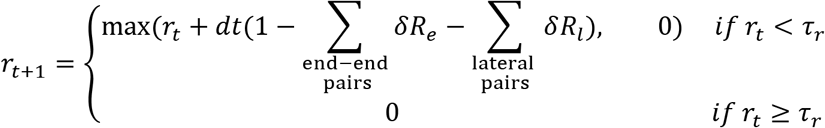

Here, *r_t_* is reversal clock at time step *t*, *r*_*t*+1_ is reversal clock at time step *t* + 1, *τ_r_* is reversal period, *δR_e_* is end-end reversal suppression factor, *δR_l_* is lateral reversal suppression factor and *dt* is the time step.

### Simulation of the mixed agent population

To simulate mixed populations of two types of agents, we assign each agent a label that corresponds to the strain it represents. We use WT label for parent strain, OE for TraAB-expression, and NR for non-reversing. Adhesion interactions are assumed to be 10x stronger (*k_adh_* = 0.1) if both agents have OE labels as compared to all other pairs (*k_adh_* = 0.01). For simulations of mixture OE agents of different TraAB alleles, no adhesion between agents with different alleles is occurring (*k_adh_* = 0), thus the reversal suppression also will not occur. Since in our model adhesion is required for reversal suppression, these interactions do not affect reversals.

### Simulation procedure

The simulation procedure here is similar to (15). We study collective behaviors of cells by simulating mechanical interactions among a large number (M) of agents on a 2D simulation region with periodic boundary conditions in an agent-based framework.

We initialize agents one by one on a square simulation region (dimension *L_sim_*) over a few initial time steps until the desired cell density (*η*) is reached. Agents are initialized in random positions over the simulation region with their orientations (*θ*) chosen randomly in the range [0, 2*π*]. Agent nodes are initialized in the straight-line configuration. During initialization, agent configurations that overlap with existing agents are rejected. After initialization, the head node for each agent is chosen between its two end-nodes with 50% probability.

At each time step of the simulation, agents move according to the various forces acting on their nodes. Changes in node positions and velocities are obtained by integrating the equations of motion based on Newton’s laws. We use the Box2D physics library (59, 60) for solving the equations of motion and for effective collision resolution. Snapshots of the simulation region, the orientation of each agent, and its node positions are recorded every minute for later analysis.

Simulations are implemented in Java programming language with a Java port of Box2D library (http://www.jbox2d.org/). The parameters of the simulation are shown in Table 2. Other parameters of the model are the same as in (15, 28). Each simulation is run for 250 min. The codes and datasets are available in the https://github.com/Igoshin-Group/CircularAggregatesPaper repository.

## Acknowledgments

We thank Beiyan Nan for FrzCD antibodies and Lee Kroos and Larry Shimkets for helpful comments. This work was supported by the National Science Foundation grants DMS-1903275 and OIS-1951025 (to OAI) and the National Institutes of Health grants R35GM140886 and GM101449 (to D.W).

## Supplemental Materials

**Fig. S1.**
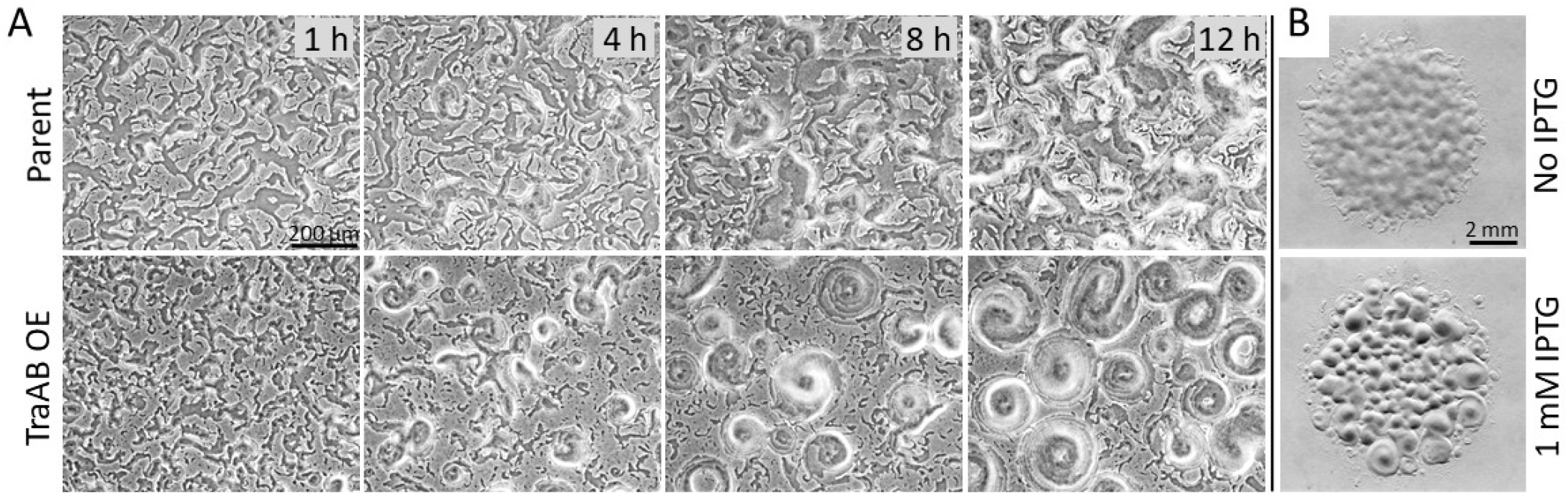
Induction of CAs. (A) Time course after plating of constitutive P*_pilA_-traAB* OE strain and its parent on agar spotted at a cell densities of 3 × 10^9^ cfu/ml. (B) Strain with an inducible promoter (P_IPTG_-*traAB* OE) plated in the presence and absence of inducer and grown for 36 h.

**Fig. S2.**
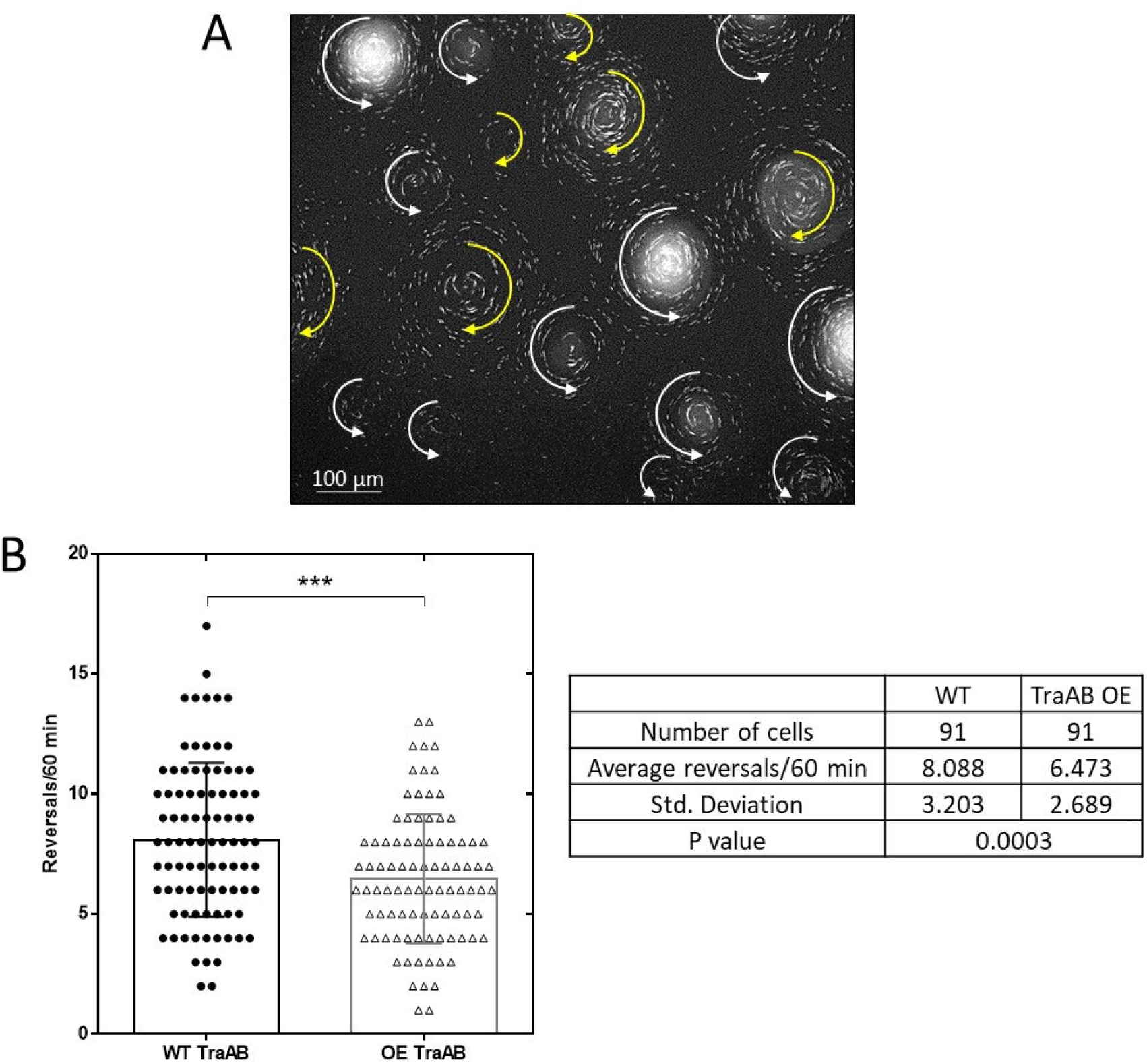
Cell tracking and reversal tracking. (A) Fluorescent micrograph of CAs. Labeled and non-labeled cells mixed 1:250. White arrows, counter-clockwise rotations; yellow arrows clockwise rotation. Time-lapse series shown in Movie S2. (B) TraAB OE does not suppress cell reversals of isolated cells at low cell density. Cell movements of isolated cells were tracked by time-lapse microscopy in three independent experiments and compared between TraAB OE to WT cells.

**Fig. S3.**
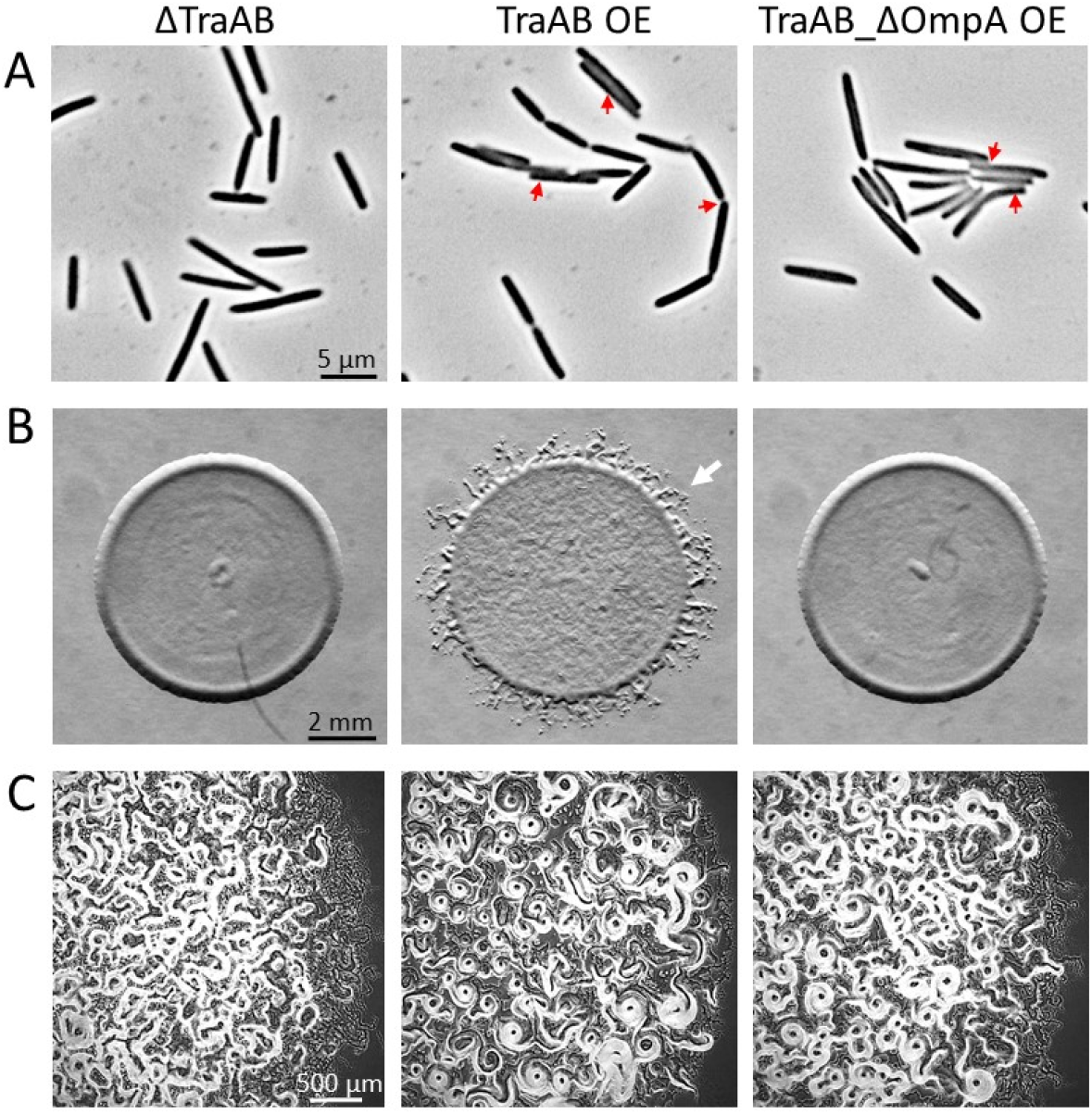
Outer membrane exchange (OME) is not required for CA formation. (A) Cell-cell adhesion absent in Δ*traAB* strain and present in two TraAB OE strains (red arrows). (B) Stimulation assays assess function of *traAB* alleles in OME. Here, two nonmotile strains were mixed 1:1 and when OME occurs the recipient cells receive missing motility lipoproteins (CglC and Tgl) and swarm out from the colony edge (middle panel arrow) (34). When OME is defective the recipient cells remain nonmotile and produce sharp colony edges (left, right panels). (C) Ability of *traAB* alleles to promote CAs (middle, right panels) or not (left panel). See Table 1 for strain details.

**Fig. S4.**
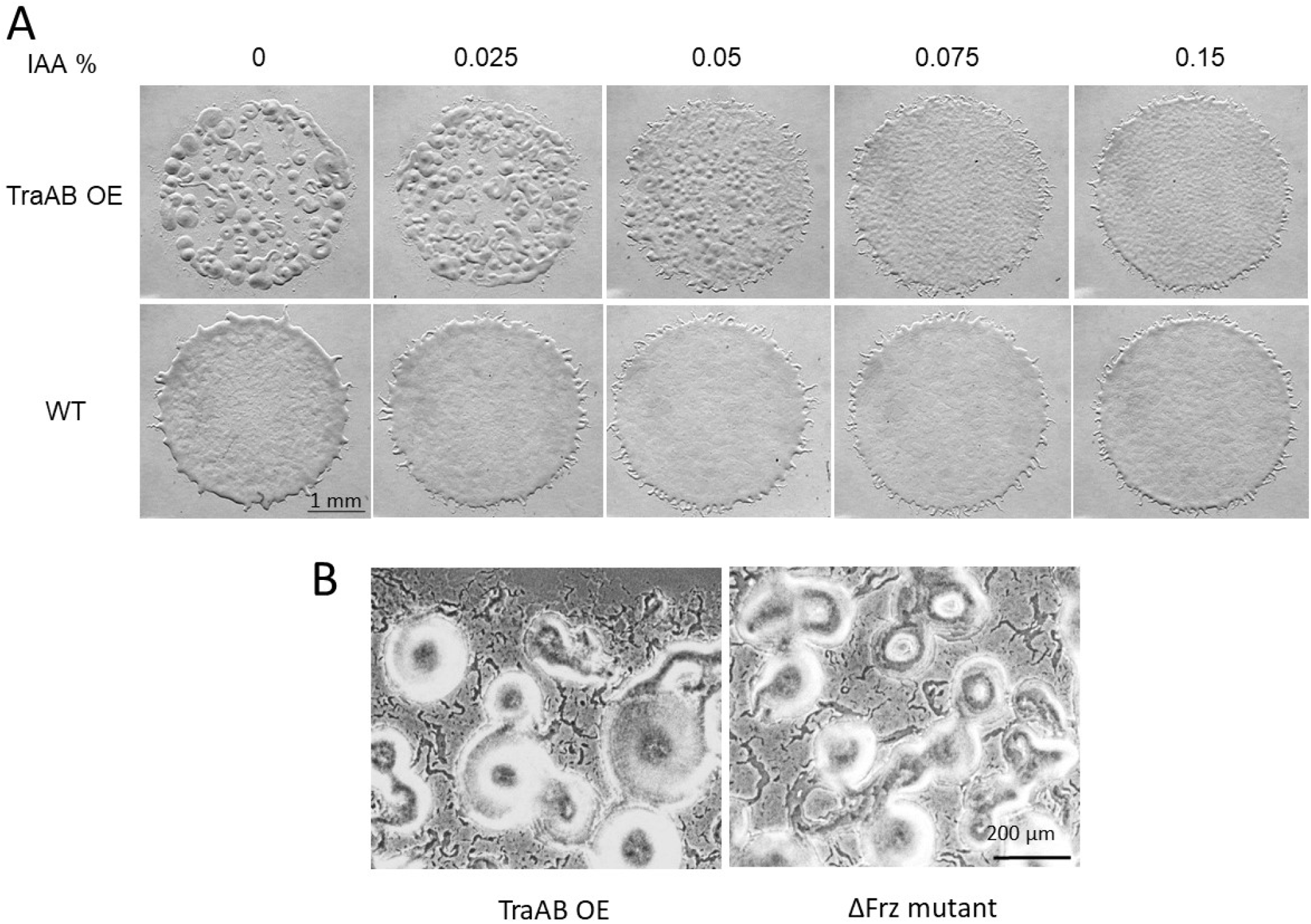
Role of Frz in CA formation. (A) Chemical induction of cellular reversals blocks CA formation. Isoamyl alcohol (IAA) stimulates cell reversals by activating the Frz pathway. IAA was added at indicated concentrations to agar media and micrographs taken at 24 h. The two left panels with no IAA are also shown in Fig. 1. (B) Frz non-reversing mutant forms CAs, though not as distinctive as the TraAB OE strain.

**Fig. S5.**
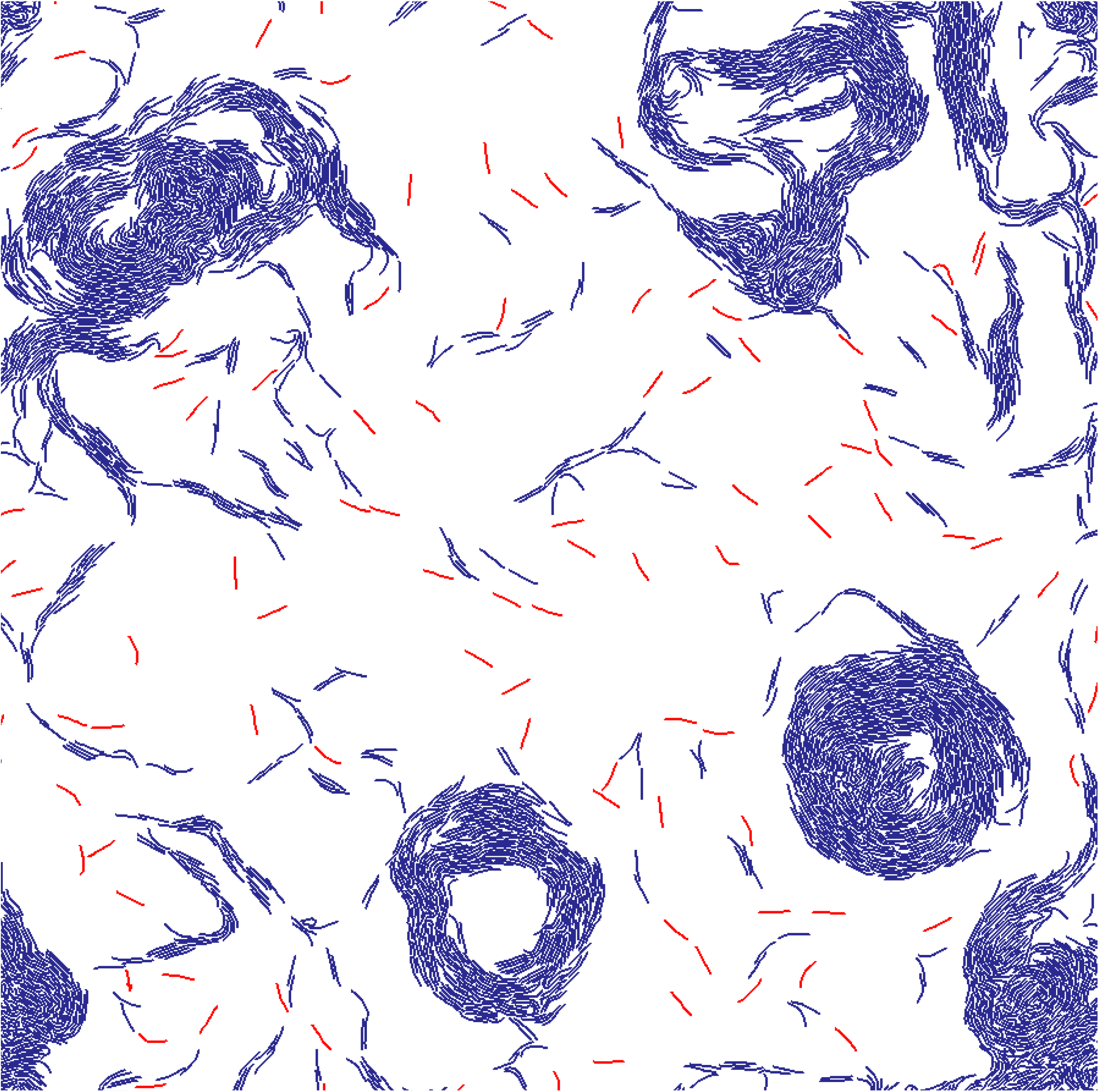
WT agents without time threshold for reversal suppression form CAs. Black agents not reversal-suppressed; blue agents are reversal-suppressed.

**Fig. S6.**
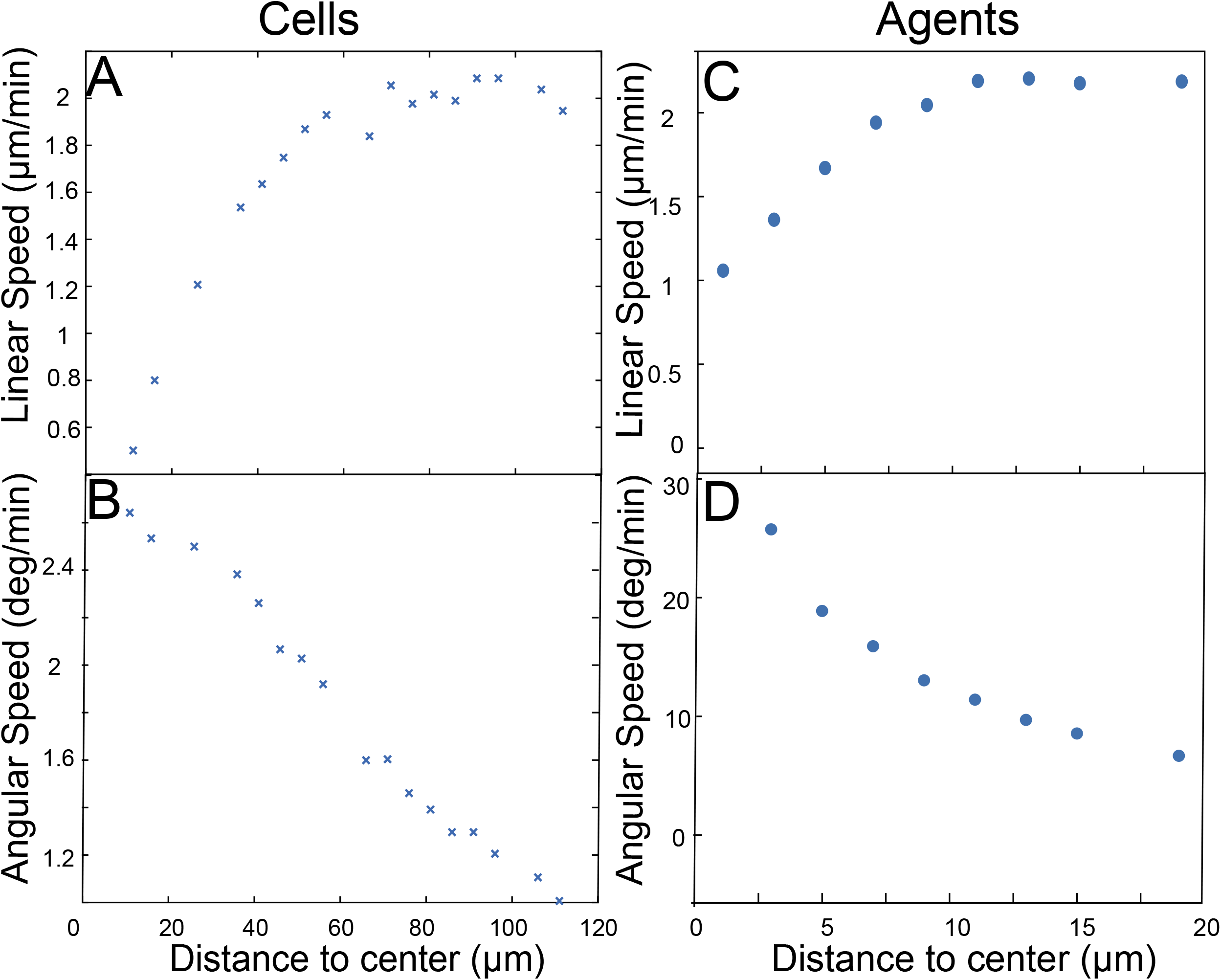
Linear speed and angular speed as measured experimentally and in simulations. Experimentally determined cell linear speed (A) and angular speed (B) measured as a function of distance to the center from Figure 3A and Movie S2. Each data point represents the average of cells within a 5 μm window. Agent linear (C) and angular speed (D) measured as a function of distance to center from simulations. Each data point represents the average of cells within a 2 μm ring. The data and simulations indicate that the CAs do not rotate as a rigid body, because that is defined as cells having the same angular speed throughout CAs. Note the quantitative agreement between the speed values is not possible because the CAs in the experiment were larger (radius 120 μm) than in the simulation (radius < 20 μm). This discrepancy was mainly due to computational limitations as our simulation can only simulate a limited number of agents in a small simulation domain (200 μm× 200 μm).

**Fig. S7.**
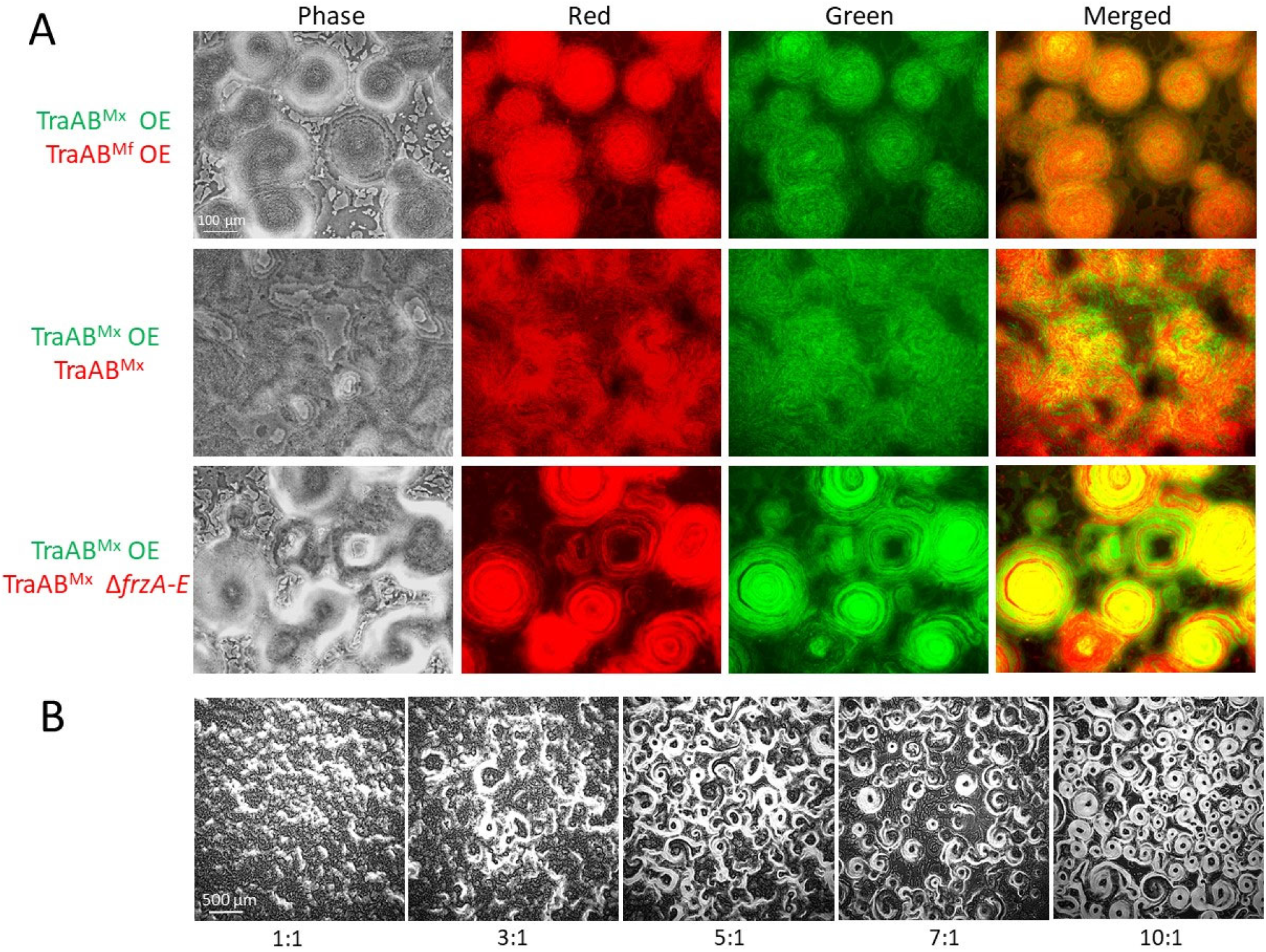
Impact of strain mixing on CA formation of TraAB OE strain. (A) CA phenotypes of strain mixtures (1:1 ratios) expressing different *traAB* alleles. Panels show single channels and merged images used to create Figure 6D-F and strains used. Media contained 1 mM IPTG to allow TraAB^Mx^ overexpression (DW2540). See Table 1 for additional strain details. (B) WT cells inhibit CA formation by TraAB OE. The ratios of TraAB OE to WT are indicated. Micrographs taken at 24 h.

**Movie S1** Time-lapse series of strains from Figure S1 at indicated times after cell plating. Time intervals between frames 30 sec; each series for 15 min. See Table 1 for strain details.

**Movie S2** Time-lapse movie that is represented in Figures 3A and S2. Time intervals between frames 1 min. See Table 1 for strain details.

**Movie S3** Time-lapse movie from Figure 4B. The movie records the last 60 mins of a 180 min simulation with a different color scheme for better contrast. In each frame, white cells represent reversal-suppressed cells and red cells are not reversal-suppressed.

## References

1. Wingreen NS, Huang KC. 2015. Physics of Intracellular Organization in Bacteria. Annu Rev Microbiol 69:361–79.

2. Couzin ID, Krause J. 2003. Self-organization and collective behavior in vertebrates, p 1-67. In Peter J.B. Slater JSR, Charles T. Snowdon, Timothy J. Roper (ed), Advances in the Study of Behavior, vol 32. Elsevier Science, San Diego, CA, USA.

3. Moussaid M, Garnier S, Theraulaz G, Helbing D. 2009. Collective information processing and pattern formation in swarms, flocks, and crowds. Top Cogn Sci 1:469–97.

4. Wall D. 2016. Kin recognition in bacteria. Annu Rev Microbiol 70:143–60.

5. Zusman DR, Scott AE, Yang Z, Kirby JR. 2007. Chemosensory pathways, motility and development in Myxococcus xanthus. Nat Rev Microbiol 5:862–72.

6. Berleman JE, Chumley T, Cheung P, Kirby JR. 2006. Rippling is a predatory behavior in Myxococcus xanthus. J Bacteriol 188:5888–95.

7. Welch R, Kaiser D. 2001. Cell behavior in traveling wave patterns of myxobacteria. Proc Natl Acad Sci U S A 98:14907–12.

8. Igoshin OA, Mogilner A, Welch RD, Kaiser D, Oster G. 2001. Pattern formation and traveling waves in myxobacteria: theory and modeling. Proc Natl Acad Sci U S A 98:14913–8.

9. Cotter CR, Schuttler HB, Igoshin OA, Shimkets LJ. 2017. Data-driven modeling reveals cell behaviors controlling self-organization during Myxococcus xanthus development. Proc Natl Acad Sci U S A 114:E4592–E4601.

10. Zhang H, Vaksman Z, Litwin DB, Shi P, Kaplan HB, Igoshin OA. 2012. The mechanistic basis of Myxococcus xanthus rippling behavior and its physiological role during predation. PLoS Comput Biol 8:e1002715.

11. Gans J, Wolinsky M, Dunbar J. 2005. Computational improvements reveal great bacterial diversity and high metal toxicity in soil. Science 309:1387–90.

12. Wielgoss S, Wolfensberger R, Sun L, Fiegna F, Velicer GJ. 2019. Social genes are selection hotspots in kin groups of a soil microbe. Science 363:1342–1345.

13. Sah GP, Wall D. 2020. Kin recognition and outer membrane exchange (OME) in myxobacteria. Curr Opin Microbiol 56:81–88.

14. Vassallo CN, Troselj V, Weltzer ML, Wall D. 2020. Rapid diversification of wild social groups driven by toxin-immunity loci on mobile genetic elements. ISME J 14:2474–2487.

15. Balagam R, Igoshin OA. 2015. Mechanism for Collective Cell Alignment in Myxococcus xanthus Bacteria. PLoS Comput Biol 11:e1004474.

16. Cao P, Wall D. 2017. Self-identity reprogrammed by a single residue switch in a cell surface receptor of a social bacterium. Proc Natl Acad Sci U S A 114:3732–3737.

17. Cao P, Wei X, Awal RP, Muller R, Wall D. 2019. A highly polymorphic receptor governs many distinct self-recognition types within the Myxococcales order. mBio 10.

18. Pathak DT, Wei X, Bucuvalas A, Haft DH, Gerloff DL, Wall D. 2012. Cell contact-dependent outer membrane exchange in myxobacteria: Genetic determinants and mechanism. PLoS Genet 8:e1002626.

19. Vassallo C, Pathak DT, Cao P, Zuckerman DM, Hoiczyk E, Wall D. 2015. Cell rejuvenation and social behaviors promoted by LPS exchange in myxobacteria. Proc Natl Acad Sci U S A 112:E2939–46.

20. Vassallo CN, Wall D. 2019. Self-identity barcodes encoded by six expansive polymorphic toxin families discriminate kin in myxobacteria. Proc Natl Acad Sci U S A 116:24808–24818.

21. Vassallo CN, Wall D. 2016. Tissue repair in myxobacteria: A cooperative strategy to heal cellular damage. BioEssays 38:306–15.

22. Nan B, Zusman DR. 2011. Uncovering the mystery of gliding motility in the myxobacteria. Annu Rev Genet 45:21–39.

23. Morrison CE, Zusman DR. 1979. Myxococcus xanthus mutants with temperature-sensitive, stage-specific defects: evidence for independent pathways in development. J Bacteriol 140:1036–42.

24. Zusman DR. 1982. “Frizzy” mutants: a new class of aggregation-defective developmental mutants of Myxococcus xanthus. J Bacteriol 150:1430–7.

25. McLoon AL, Wuichet K, Hasler M, Keilberg D, Szadkowski D, Sogaard-Andersen L. 2016. MglC, a Paralog of Myxococcus xanthus GTPase-Activating Protein MglB, Plays a Divergent Role in Motility Regulation. J Bacteriol 198:510–20.

26. Starruss J, Peruani F, Jakovljevic V, Sogaard-Andersen L, Deutsch A, Bar M. 2012. Pattern-formation mechanisms in motility mutants of Myxococcus xanthus. Interface Focus 2:774–85.

27. Jelsbak L, Sogaard-Andersen L. 2000. Pattern formation: fruiting body morphogenesis in Myxococcus xanthus. Curr Opin Microbiol 3:637–42.

28. Balagam R, Litwin DB, Czerwinski F, Sun M, Kaplan HB, Shaevitz JW, Igoshin OA. 2014. Myxococcus xanthus gliding motors are elastically coupled to the substrate as predicted by the focal adhesion model of gliding motility. PLoS Comput Biol 10:e1003619.

29. Burchard RP. 1982. Trail following by gliding bacteria. J Bacteriol 152:495–501.

30. Ducret A, Valignat MP, Mouhamar F, Mignot T, Theodoly O. 2012. Wet-surface-enhanced ellipsometric contrast microscopy identifies slime as a major adhesion factor during bacterial surface motility. Proc Natl Acad Sci U S A 109:10036–41.

31. Wolgemuth C, Hoiczyk E, Kaiser D, Oster G. 2002. How myxobacteria glide. Curr Biol 12:36977.

32. Perez-Burgos M, Sogaard-Andersen L. 2020. Biosynthesis and function of cell-surface polysaccharides in the social bacterium Myxococcus xanthus. Biol Chem 401:1375–1387.

33. Gloag ES, Turnbull L, Javed MA, Wang H, Gee ML, Wade SA, Whitchurch CB. 2016. Stigmergy co-ordinates multicellular collective behaviours during Myxococcus xanthus surface migration. Sci Rep 6:26005.

34. Pathak DT, Wall D. 2012. Identification of the *cglC*, *cglD*, *cglE*, and *cglF* genes and their role in cell contact-dependent gliding motility in *Myxococcus xanthus*. J Bacteriol 194:1940–9.

35. Bustamante VH, Martinez-Flores I, Vlamakis HC, Zusman DR. 2004. Analysis of the Frz signal transduction system of Myxococcus xanthus shows the importance of the conserved C-terminal region of the cytoplasmic chemoreceptor FrzCD in sensing signals. Mol Microbiol 53:1501–13.

36. McBride MJ, Kohler T, Zusman DR. 1992. Methylation of FrzCD, a methyl-accepting taxis protein of Myxococcus xanthus, is correlated with factors affecting cell behavior. J Bacteriol 174:4246–57.

37. Herrou J, Mignot T. 2020. Dynamic polarity control by a tunable protein oscillator in bacteria. Curr Opin Cell Biol 62:54–60.

38. Pogue CB, Zhou T, Nan B. 2018. PlpA, a PilZ-like protein, regulates directed motility of the bacterium Myxococcus xanthus. Mol Microbiol 107:214–228.

39. Parkinson JS, Hazelbauer GL, Falke JJ. 2015. Signaling and sensory adaptation in Escherichia coli chemoreceptors: 2015 update. Trends Microbiol 23:257–66.

40. Collins KD, Lacal J, Ottemann KM. 2014. Internal sense of direction: sensing and signaling from cytoplasmic chemoreceptors. Microbiol Mol Biol Rev 78:672–84.

41. Xu Q, Black WP, Cadieux CL, Yang Z. 2008. Independence and interdependence of Dif and Frz chemosensory pathways in Myxococcus xanthus chemotaxis. Mol Microbiol 69:714–23.

42. McCleary WR, McBride MJ, Zusman DR. 1990. Developmental sensory transduction in Myxococcus xanthus involves methylation and demethylation of FrzCD. J Bacteriol 172:4877–87.

43. Jelsbak L, Sogaard-Andersen L. 2002. Pattern formation by a cell surface-associated morphogen in Myxococcus xanthus. Proc Natl Acad Sci U S A 99:2032–7.

44. Shi W, Ngok FK, Zusman DR. 1996. Cell density regulates cellular reversal frequency in Myxococcus xanthus. Proc Natl Acad Sci U S A 93:4142–6.

45. Janulevicius A, Van Loosdrecht M, Picioreanu C. 2015. Short-Range Guiding Can Result in the Formation of Circular Aggregates in Myxobacteria Populations. PLOS Computational Biology 11:e1004213.

46. Vassallo CN, Cao P, Conklin A, Finkelstein H, Hayes CS, Wall D. 2017. Infectious polymorphic toxins delivered by outer membrane exchange discriminate kin in myxobacteria. eLife 6.

47. Kroos L. 2017. Highly Signal-Responsive Gene Regulatory Network Governing Myxococcus Development. Trends Genet 33:3–15.

48. Boynton TO, Shimkets LJ. 2015. Myxococcus CsgA, Drosophila Sniffer, and human HSD10 are cardiolipin phospholipases. Genes Dev 29:1903–14.

49. Li S, Lee BU, Shimkets LJ. 1992. csgA expression entrains Myxococcus xanthus development. Genes Dev 6:401–10.

50. Lobedanz S, Sogaard-Andersen L. 2003. Identification of the C-signal, a contact-dependent morphogen coordinating multiple developmental responses in Myxococcus xanthus. Genes Dev 17:2151–61.

51. Schumacher D, Sogaard-Andersen L. 2017. Regulation of Cell Polarity in Motility and Cell Division in Myxococcus xanthus. Annu Rev Microbiol 71:61–78.

52. Bonner PJ, Xu Q, Black WP, Li Z, Yang Z, Shimkets LJ. 2005. The Dif chemosensory pathway is directly involved in phosphatidylethanolamine sensory transduction in Myxococcus xanthus. Mol Microbiol 57:1499–508.

53. Kuzmich S, Skotnicka D, Szadkowski D, Klos P, Perez-Burgos M, Schander E, Schumacher D, Sogaard-Andersen L. 2021. Three PilZ Domain Proteins, PlpA, PixA, and PixB, Have Distinct Functions in Regulation of Motility and Development in Myxococcus xanthus. J Bacteriol 203:e0012621.

54. Zhou T, Nan B. 2017. Exopolysaccharides promote Myxococcus xanthus social motility by inhibiting cellular reversals. Mol Microbiol 103:729–743.

55. Iniesta AA, Garcia-Heras F, Abellon-Ruiz J, Gallego-Garcia A, Elias-Arnanz M. 2012. Two systems for conditional gene expression in Myxococcus xanthus inducible by isopropyl-beta-D-thiogalactopyranoside or vanillate. J Bacteriol 194:5875–85.

56. Sun M, Wartel M, Cascales E, Shaevitz JW, Mignot T. 2011. Motor-driven intracellular transport powers bacterial gliding motility. Proceedings of the National Academy of Sciences 108:7559–7564.

57. Mignot T, Shaevitz JW, Hartzell PL, Zusman DR. 2007. Evidence that focal adhesion complexes power bacterial gliding motility. Science 315:853–6.

58. Sliusarenko O, Zusman DR, Oster G. 2007. The motors powering A-motility in Myxococcus xanthus are distributed along the cell body. J Bacteriol 189:7920–1.

59. Pugsley AP. 1992. Translocation of a folded protein across the outer membrane in Escherichia coli. Proc Natl Acad Sci U S A 89:12058–62.

60. Catto E. 2012. Box2D - A 2D physics engine for games. https://box2d.org/. Accessed

61. Pathak DT, Wei X, Dey A, Wall D. 2013. Molecular recognition by a polymorphic cell surface receptor governs cooperative behaviors in bacteria. PLoS Genet 9:e1003891.

62. Dey A, Vassallo CN, Conklin AC, Pathak DT, Troselj V, Wall D. 2016. Sibling rivalry in *Myxococcus xanthus* is mediated by kin recognition and a polyploid prophage. J Bacteriol 198:994–1004.

63. Wall D, Kaiser D. 1998. Alignment enhances the cell-to-cell transfer of pilus phenotype. Proc Natl Acad Sci U S A 95:3054–8.

64. Wu SS, Kaiser D. 1997. Regulation of expression of the pilA gene in Myxococcus xanthus. J Bacteriol 179:7748–58.

65. Janulevicius A, Van Loosdrecht MCM, Simone A, Picioreanu C. 2010. Cell Flexibility Affects the Alignment of Model Myxobacteria. Biophysical Journal 99:3129–3138.

66. Harvey CW, Morcos F, Sweet CR, Kaiser D, Chatterjee S, Liu X, Chen DZ, Alber M. 2011. Study of elastic collisions of Myxococcus xanthus in swarms. Phys Biol 8:026016.

67. Wolgemuth CW. 2005. Force and Flexibility of Flailing Myxobacteria. Biophysical Journal 89:945–950.

